# Right posterior theta reflects human parahippocampal phase resetting by salient cues during goal-directed navigation

**DOI:** 10.1101/2024.11.16.623803

**Authors:** Malte R. Güth, Andrew Reid, Yu Zhang, Sonja C. Huntgeburth, Ravi D. Mill, Alain Dagher, Kim Kerns, Clay B. Holroyd, Michael Petrides, Michael W. Cole, Travis E. Baker

**Author notes:** Correspondence:* Malte R. Güth, 312 Church St. SE, Minneapolis, MN 55455.

## Abstract

Animal and computational work indicate that phase resetting of theta oscillations (4-12 Hz) in the parahippocampal gyrus (PHG) by salient events (e.g., reward, landmarks) facilitates the encoding of goal-oriented information during navigation. Although well-studied in animals, this mechanism has not been empirically substantiated in humans. In the present article, we present data from two studies (Study 1: asynchronous EEG-MEG | Study 2: simultaneous EEG-fMRI) to investigate theta phase resetting and its relationship to PHG BOLD activation in healthy adults (aged 18-34 years old) navigating a virtual T-maze to find rewards. In the first experiment, both EEG and MEG data revealed a burst of theta power over right-posterior scalp locations following feedback onset (termed right-posterior theta, RPT), and RPT power and measures of phase resetting were sensitive to the subject’s spatial trajectory. In Experiment 2, we used probabilistic tractography data from the human connectome project to segment the anterior and posterior PHG based on differential connectivity profiles to other brain regions. This analysis resulted in a PHG subdivision consisting of four distinct anterior and two posterior PHG clusters. Next, a series of linear mixed effects models based on simultaneous EEG-fMRI data revealed that single-trial RPT peak power significantly predicted single-trial hemodynamic responses in two clusters within the posterior PHG and one in the anterior PHG. This coupling between RPT power and PHG BOLD was exclusive to trials performed during maze navigation, and not during a similar task devoid of the spatial context of the maze. These findings highlight a role of PHG theta phase resetting for the purpose of encoding salient information during goal-directed spatial navigation. Taken together, RPT during virtual navigation integrates experimental, computational, and theoretical research of PHG function in animals with human cognitive electrophysiology studies and clinical research on memory-related disorders such as Alzheimer’s disease.

## Introduction

Finding refuge, food, or mates depends on an animal’s ability to accurately encode salient events within the context of their goals during navigation. Prior theoretical and animal work indicate that the timing of neuronal activity within the hippocampus and parahippocampal gyrus (PHG; in rodents perirhinal, postrhinal, and entorhinal cortical areas) with respect to the ongoing local field potential oscillating at a theta rhythm (4-12 Hz in rodents) plays a critical role in this cortical process (Hasselmo et al., 2009; Mizuseki et al., 2009; Newman & Hasselmo, 2014). Notably, when salient events or cues such as rewards and navigationally-relevant landmarks are presented in the animal’s environment, the phase of the theta rhythm is reset (Buzsáki et al., 1979; Givens, 1996; Williams & Givens, 2003), a process that appears to enhance event encoding (Hasselmo, 2008; Hyman et al., 2003; McCartney et al., 2004; Vinogradova et al., 1996). Animal and computational work indicate that event encoding critically depends on the phase, frequency, and temporal dynamics of PHG theta oscillations (Hasselmo & Stern, 2014; Hölscher et al., 1997; Newman & Hasselmo, 2014; Quirk et al., 2021). For example, it has been argued that event-locked reset of the ongoing theta local field potential aligns imperative information with optimal conditions for event encoding and long-term potentiation (Hasselmo, 2008; Hasselmo et al., 2009). Computational models of spatial navigation make use of this mechanism to simulate PHG memory encoding and retrieval of events encountered during spatial navigation (Hasselmo, 2008; Hasselmo et al., 2009). In particular, phase resetting of PHG theta oscillations by location-specific input from place cells is thought to prevent accumulation of error, facilitate the encoding of landmarks and goal locations, and to enhance path integration during navigation (Burgess, 2008; Burgess et al., 2007; Hasselmo, 2008). Although phase resetting of PHG theta oscillations during navigation has been previously studied in animal and computational work, this mechanism remains poorly understood in the human brain because of the limitations of non-invasive neuroimaging methods. Comparing rodent and human research is complicated by the anatomical differences in PHG definitions (in humans perihinal, parahippocampal, entorhinal cortical areas; Burwell, 2000; Burwell et al., 1995) and a lack of consensus on human PHG delineation (Syversen et al., 2021).

Over the past decades, intracranial electrophysiological recordings in epilepsy patients have demonstrated the presence of movement-related theta oscillations in both the neocortex (Kahana et al., 1999) and hippocampus (Ekstrom et al., 2005) during immobile virtual navigation.

Electroencephalography (EEG) and magnetoencephalography (MEG) studies have also identified functional parallels between theta oscillations found in rodents during active navigation and those recorded in humans (4-8 Hz) during virtual navigation using tasks involving spatial learning and spatial memory (Cornwell et al., 2008; Jacobs et al., 2006), self-initiated movement, speed, and direction (M.-H. Lin et al., 2022), processing of landmarks (Cornwell et al., 2008; Rounds et al., 2020), as well as path integration and orientation (C.-T. Lin et al., 2015). Recent non-human primate intracranial (Mao, 2023; Mao et al., 2021) and human functional magnetic resonance imaging (fMRI) studies are also suggestive of the presence of theta phase coding. For instance, Doeller et al. (2010) observed that in a foraging task the fMRI blood-oxygenation-level-dependent (BOLD) response in the right human PHG showed a periodic increase of BOLD signal in 60 degree increments of angular running direction (i.e., six-fold rotational symmetry) consistent with the predictions of theoretical models of theta phase coding (Burgess, 2008; Burgess et al., 2007). Together, these findings support the idea that this area in human brains constitutes a “Parahippocampal Place Area” or the “Parahippocampal Spatial Scene Area” (Epstein et al., 1999, 2017; Epstein & Kanwisher, 1998), exhibiting a similar theta phase coding as reported in rodents. As such, it would be a likely source of phase resetting in response to the encoding of salient information during spatial navigation in humans.

Of particular relevance, a series of EEG studies demonstrated that feedback cues presented in a virtual T-maze environment elicit a burst of theta oscillations (4-12 Hz) over posterior electrode locations, with larger amplitudes over right hemisphere channels (Baker & Holroyd, 2009, 2013). This right posterior theta (RPT) response (Figure 1) was characterized by spectral power peaking in the theta band around 200 ms following feedback onset, and was consistent with a partial phase reset of the oscillation (i.e., concomitant increases in phase coherence and spectral power relative to baseline; Min et al., 2007; Yeung et al., 2004). We proposed that RPT arises from phase resetting and enhancement in power of ongoing EEG theta activity: RPT is generated when an event leads to resetting of the phase of ongoing EEG oscillations, such that peaks and troughs in the oscillatory waveform become aligned – that is, time-locked to the resetting event. In addition, RPT power enhancement captures the increase in PHG theta power at the time of event encoding that has been previously observed intracranially (e.g., Lega et al., 2012; Miller et al., 2018). Further, the latency, power, and phase angle of RPT was found to be sensitive to feedback cues received following a right turn relative to left turn in the T-maze (Baker & Holroyd, 2013). RPT peaked earlier with greater power for right alleys, which was accompanied by higher alignment of phase angles. EEG source localization analysis and fMRI data point to the PHG as a likely source of RPT (Baker et al., 2015; Baker & Holroyd, 2013). In particular, our previous fMRI study demonstrated that when comparing feedback processing in the T-maze task to the same task devoid of the spatial context (i.e., No-maze), there was an increase in the hemodynamic response in the PHG, medial temporal-occipital cortex, and right precuneus, regions commonly activated during virtual navigation (Baker et al., 2015). Moreover, posterior and anterior PHG regions were more activated by feedback stimuli following right compared to left turns only when navigating in the maze. Both regions also exhibited heightened activation in individuals with greater spatial ability (Baker et al., 2015), findings consistent with EEG data (Baker & Holroyd, 2013). Together, this literature supports the hypothesis that the right PHG provides the neural substrate for the encoding of salient events within the larger system for navigation, that the PHG theta rhythm is anchored to salient events via phase resetting, and that this reset impacts encoding by shifting the timing of neuronal firing in relation to the phase of the theta rhythm (Aguirre et al., 1996; Baker et al., 2015; Ekstrom et al., 2003; Maguire, Frith, et al., 1998).

**Figure 1.**
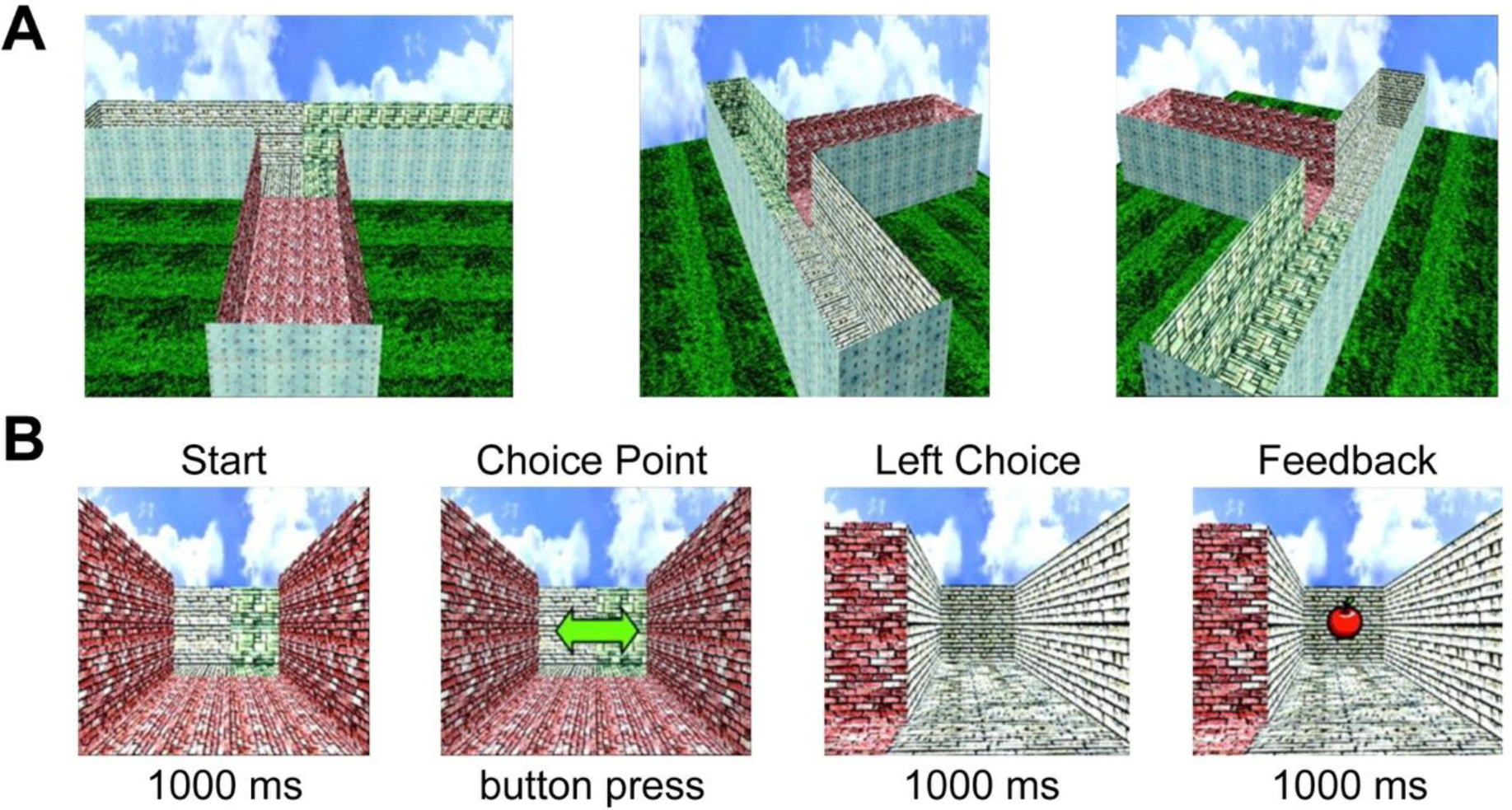
Overview of the virtual T-maze task. A: Shape of the virtual T-maze as seen from above focused on the start position (left), the left alley (center) and the right alley (right). B: Sequence of events comprising an example trial of the T-maze task. Top line is the event label and bottom line shows the event duration. Choice point image lasts until subject performs a button press. Behavioral analyses testing the equivalence of left and right alleys (i.e., reaction times, choice frequency) and choice patterns are provided in detail in Figure 1-figure supplement 1.

Based on the results of the studies described above, we proposed that reward-related stimuli in the maze induced a partial phase reset of the PHG theta rhythm with relatively lower latency, greater power, and greater phase alignment in right alleys compared to left alleys in the maze, which we term here the RPT effect. This phenomenon can also be observed in the time domain as a component of the event-related brain potential (ERP) called the topographical N170 (Baker & Holroyd, 2009, 2013). However, this proposal is complicated by two factors. First, electrical fields are distorted by varying conductivities of tissues between the source and the electrode (i.e., volume conduction), which can in turn distort the timing and power of the recorded signal between experimental conditions (Nunez, 1981; Nunez et al., 1997; Rush & Driscoll, 1969). Second, while source localization analyses have shown that the PHG is a likely source of RPT, they do not allow for any conclusions about the temporal covariation between PHG activity and RPT across trials. Furthermore, as the exact relation between the fMRI BOLD responses and underlying oscillatory activity is not yet fully understood (Ekstrom, 2010, 2021; Logothetis, 2008; Logothetis & Wandell, 2004), it remains to be explored whether the BOLD response observed in our previous fMRI study was indeed related to RPT.

To overcome these limiting factors, we first examined whether the RPT effect was reproducible using EEG and MEG data recorded from the same individuals. While EEG allows for the recording of activity from dipoles of varying orientations and depths (Ahlfors et al., 2010; D. Cohen & Cuffin, 1991), MEG allows for the recording of activity from tangentially oriented dipoles at a high signal-to-noise ratio (Baillet et al., 2001; Melcher & Cohen, 1988), with less distortion of oscillations through volume conduction than EEG (van den Broek et al., 1998). Replicating the RPT effect using MEG would help rule out potential confounding sources of the RPT effect. Second, we set out to examine the relationship between the BOLD signal and RPT power changes by simultaneously recording EEG and fMRI while human subjects engaged in both a spatial and a non-spatial navigation task to find rewards. These experiments aim to provide converging evidence for the proposal that RPT is produced by a PHG system specific to spatial navigation, thereby integrating experimental, computational, and theoretical findings from studies of animal hippocampal-parahippocampal function with those from the field of human EEG research.

## Results

### Experiment 1: Asynchronous EEG-MEG

The purpose of Experiment 1 was to use EEG (session 1) and MEG (session 2) data to test whether reward-related RPT reflects phase resetting of the ongoing theta rhythm and is sensitive to the spatial context of the feedback in the T-maze (i.e., replicate the findings from Baker and Holroyd, 2013 using EEG and MEG data collected from the same individuals). In the virtual T-maze task (Figure 1), subjects (N = 11; MAge = 25 ± 2.9 years; 5 female) navigated towards a left or right alley in a T-maze and were asked to maximize the amount of money earned by finding as many apples (5 cents per apple) as possible. Based on our previous findings, we predicted that (i) time-frequency analysis should reveal similar feedback-related RPT and phase resetting dynamics across both EEG and MEG datasets, and (ii) the RPT power, RPT latency, and phase resetting should be sensitive to the spatial location of the feedback received. As a partial phase reset is an abrupt shift in the ongoing phase of an oscillation, phase angles at the timing of RPT should align and increase in consistency across trials, as indexed by inter-trial coherence (ITC) and an analysis of the variability of phase angles.

### The RPT effect is observable in EEG and MEG

Figure 2 illustrates the results of the time-frequency analysis of the EEG and MEG response to feedback encountered in the left and right alleys (EEG left panels, MEG right panels). Visual inspection of Figure 2A and 2B reveals that for both EEG and MEG data, there was a clear enhancement of theta power (4-12 Hz) peaking approximately 100-250 ms following the onset of the feedback stimulus in both the left and right alleys. This increase in power exhibited a right posterior scalp distribution with a maximum at channel location PO8 for EEG, and right parietotemporal sensor (MRO33) for MEG (see Figure S2-figure supplement 1A). Regarding EEG activity recorded at PO8, RPT power peaked earlier for feedback found in the right alley (M = 179.63 ms, SD = ±33.62 ms) compared to feedback found in the left alley (M = 195.64 ms, SD = ±26.28 ms), *W* = 0, *p* = 0.007, *d* = 0.51 (Figure 2C), replicating our previous findings (Baker & Holroyd, 2013). This RPT latency effect of approximately 16 ms (SD = ±17.48 ms) was not observed at the contralateral electrode channel (PO7), *W* = 27.5, *p* = 1, *d* = 0.03, indicating an asymmetric spatial sensitivity of the right hemisphere. With regard to peak power, RPT power was larger for feedback found in the right alley (M = 1.24; SD = ±1.21) compared to the left alley (M = 1; SD = ±1.03), *W* = 2, *p* = 0.015, *d* = 0.2, consistent with previous findings (Baker & Holroyd, 2013). Again, this effect was not observed at the contralateral electrode channel (PO7), *W* = 17, *p* = 0.285, *d* = 0.08 (see Figure S2-figure supplement 1B).

**Figure 2.**
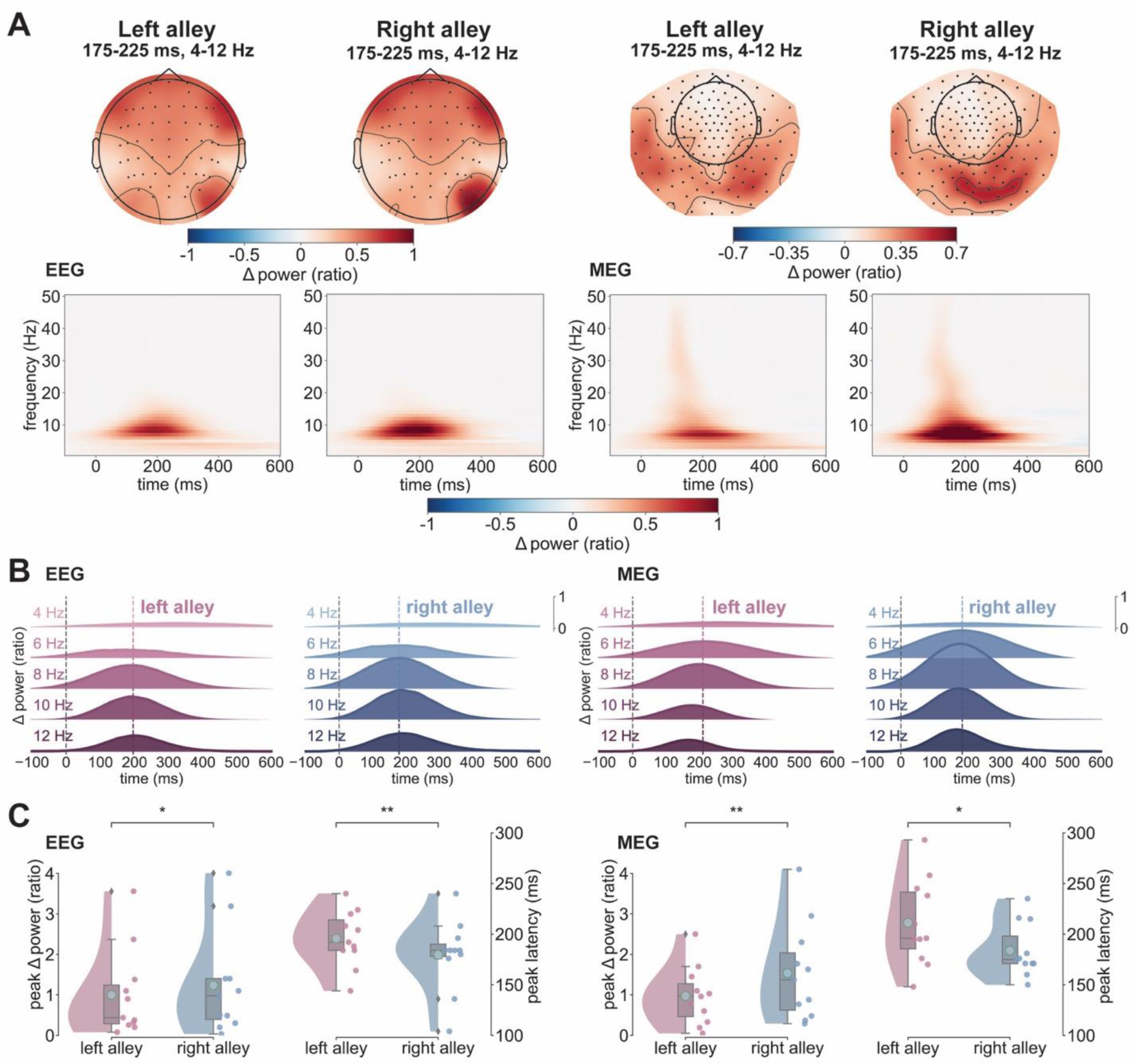
RPT effect in EEG and MEG. **A**: Spectrograms with evoked power change relative to baseline (expressed as ratio) from -100 ms to 600 ms post-feedback separated by alleys (left vs. right alley) and modality (EEG and MEG). Spectral power is unitless because it is calculated as a proportional increase/decrease relative to baseline. Topographies at the top depict the average spatial distribution of theta power from 175 ms to 225 ms. The left side of the panel contains the results for the EEG and the right side for the MEG. **B**: Ridge plots depicting average power change for individual frequency bands (1-4 Hz, 5-6 Hz, 7-8 Hz, 9-10 Hz, 11-12 Hz) between -100 ms and 600 ms post-feedback separated by left (in pink) and right alleys (in blue) and modality (EEG left side, MEG right side). Dotted colored lines mark the average peak latency across subjects for the respective alley and modality. **C**: Raincloud plots showing the distributions of theta power peaks and theta power peak latencies by alley (left in pink, right in blue) and modality (EEG on the left, MEG on the right). Cyan dots mark the mean value of each distribution and gray diamonds mark outlier values. Red and blue scatterplot dots represent individual subject values. *\*p < 0.05, **p < 0.01*

A comparable analysis on MEG recordings revealed an elevated RPT response extending from medial occipital to right parietotemporal sensors, peaking on average around 196 ms (SD = 28.59 ms; Figure 2A and 2B). For the MEG data, we identified the channels of interest by identifying the maximal peak power following the onset of feedback at each channel (averaged across conditions) and frequency. This analysis revealed a cohesive cluster of right parieto- occipital sensors displaying the largest increase in theta power following feedback onset, which were pooled together for all subsequent analysis (Figure 2A and 2B, right panels). Consistent with the EEG data, RPT power peaked earlier for feedback found in the right alley (M = 183.72 ms, SD = ±25.19) compared to the left alley (M = 211.27 ms, SD = ±40.93 ms), *W* = 8.5, *p* = 0.029, *d* = 0.79 (Figure 2C). Further, RPT exhibited a greater increase in power for right alleys (M = 1.54; SD = ±1.14) compared to left alleys (M = 0.97; SD = ±0.69), *W* = 4, *p* = 0.009, *d* = 0.57. Underscoring the connection between the RPT effect found in EEG and these MEG results was a significant Pearson correlation between RPT amplitude for right alleys across subjects (*r*(10) = 0.616, *p* = 0.044).

### Direction modulates phase coherence enhancement

Figure 3A presents an example of the single-trial EEG epochs associated with feedback onset, recorded at EEG channel PO8 (Figure 3A left panel, EEG) and the right parietooccipital MEG sensor MRO21 (Figure 3A right panel, MEG) for a single subject. The data are sorted by phase (−π to π, top to bottom) relative to feedback onset for feedback stimuli encountered in the left (top) and right (bottom) alleys. Because we had previously identified RPT bands around 8 Hz and 10 Hz (2 Hz bandwidth; 7-10 Hz) as the frequencies with the strongest phase consistency and highest sensitivity to spatial context (left vs. right alley) (Baker & Holroyd, 2013), all subsequent analyses focused on these high theta frequency bands. Note that the EEG and MEG data are color-coded such that extreme negative voltages are plotted in blue and extreme positive voltages are plotted in red. Visual inspection of Figure 3A shows strong phase consistency across trials for this subject, an impression confirmed by an analysis of ITC values, a measure of the degree to which phases of a particular frequency range synchronize across trials. Strong ITC is consistent with (Makeig et al., 2002) but not diagnostic of (Yeung et al., 2004, 2007) phase resetting of EEG oscillations at that frequency band, since strong ITC can occur without a reset.

**Figure 3.**
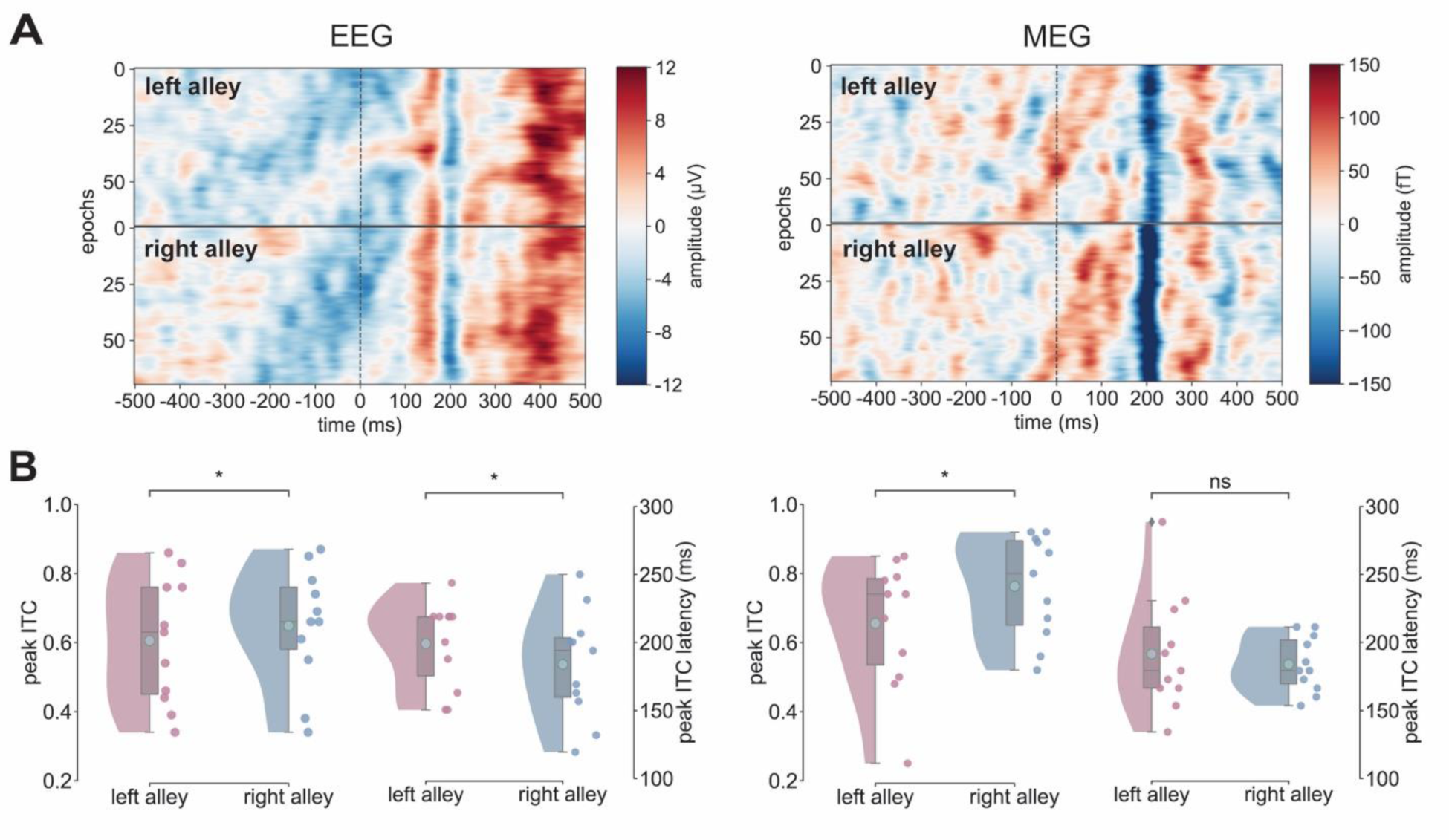
ITC magnitude and latency in EEG and MEG varies by direction. **A**: First 70 phase- ordered time-domain trials of subject 8 separated by modality (EEG on the left, MEG on the right) and alley (left in the top, right in the bottom) time-locked to feedback presentation. Color-coding represents amplitude (i.e., µV and fT) across time (-500 ms to 500 ms). **B**: Peak ITC values and peak timings (EEG left two panels, MEG right two panels) with individual subject values plotted as colored dots separated by alley (left vs. right). ns: not significant *\*p < 0.05*

Regarding EEG recordings, an analysis of phase coherence in the high theta band (7-10 Hz) revealed a large increase in ITC (M = 0.63 μV, SD = 0.16 μV) peaking around 191 ms (SD = 32.96 ms) following feedback onset. This ITC peak increase was significantly larger for right alleys (M = 0.65 μV, SD = 0.16 μV) than for left alleys (M = 0. 61 μV, SD = 0.17 μV), *W* = 12, *p* = 0.031, *d* = 0.24. In addition, the peak timing of ITC was significantly earlier for right alleys (M = 183.84 ms, SD = 38.18 ms) than for left alleys (M = 199.11 ms, SD = 30.33 ms), *W* = 49, *p* = 0.014, *d* = 0.42 (Figure 3B). Both results are a replication of our precious findings (Baker & Holroyd, 2013). In regard to the MEG data, an analysis of phase coherence in the high theta band (7-10 Hz) also revealed a large increase in ITC (M = 0.71 fT, SD = 0.15 fT) peaking around 188 ms (SD = 22.38 ms) following feedback onset. The peak ITC value and their latencies exhibited the same pattern of results as in the EEG. Right alleys were marked by a significantly larger ITC peak (M = 0.76 fT, SD = 0.14 fT) than left alleys (M = 0.66 fT, SD = 0.18 fT; *W* = 9, *p* = 0.016, *d* = 0.64), but there were no differences between left (M = 191.52 ms, SD = 41.1 ms) and right alleys (M = 188.92 ms, SD = 19.41 ms) in terms of ITC peak latencies, *W* = 28, *p* = 0.52, *d* = 0.23.

### Salient events align EEG and MEG phases across trials

Figure 4A shows circular histograms of the phase distribution for the frequency band associated with maximal EEG-related (left panels) and MEG-related (right panels) RPT power relative to pre- and post-stimulus onset for a representative subject. Figure 4B shows the resultant vector length (RVL) of phase angles across subjects for the right (red) and left (blue) alleys recorded at channel PO8 for EEG and right parietotemporal sensor for MEG. Following the ITC analysis, the phase analysis was focused on high theta frequency bands (7-10 Hz). To test whether the spread of phase angles and thus the degree of phase alignment was different between pre- and post-feedback windows, an initial Wilcoxon signed-rank test was run to compare the RVL mean during the pre-feedback window to the post-feedback window. This was then followed by a two-factor Harrison-Kanji test on the circular grand mean (CGM) values with Feedback Alley Type (left vs. right alley), and Frequency (8 Hz vs. 10 Hz bands) as factors. For the RVL the same factor model was implemented a standard two-way repeated measures ANOVA.

**Figure 4.**
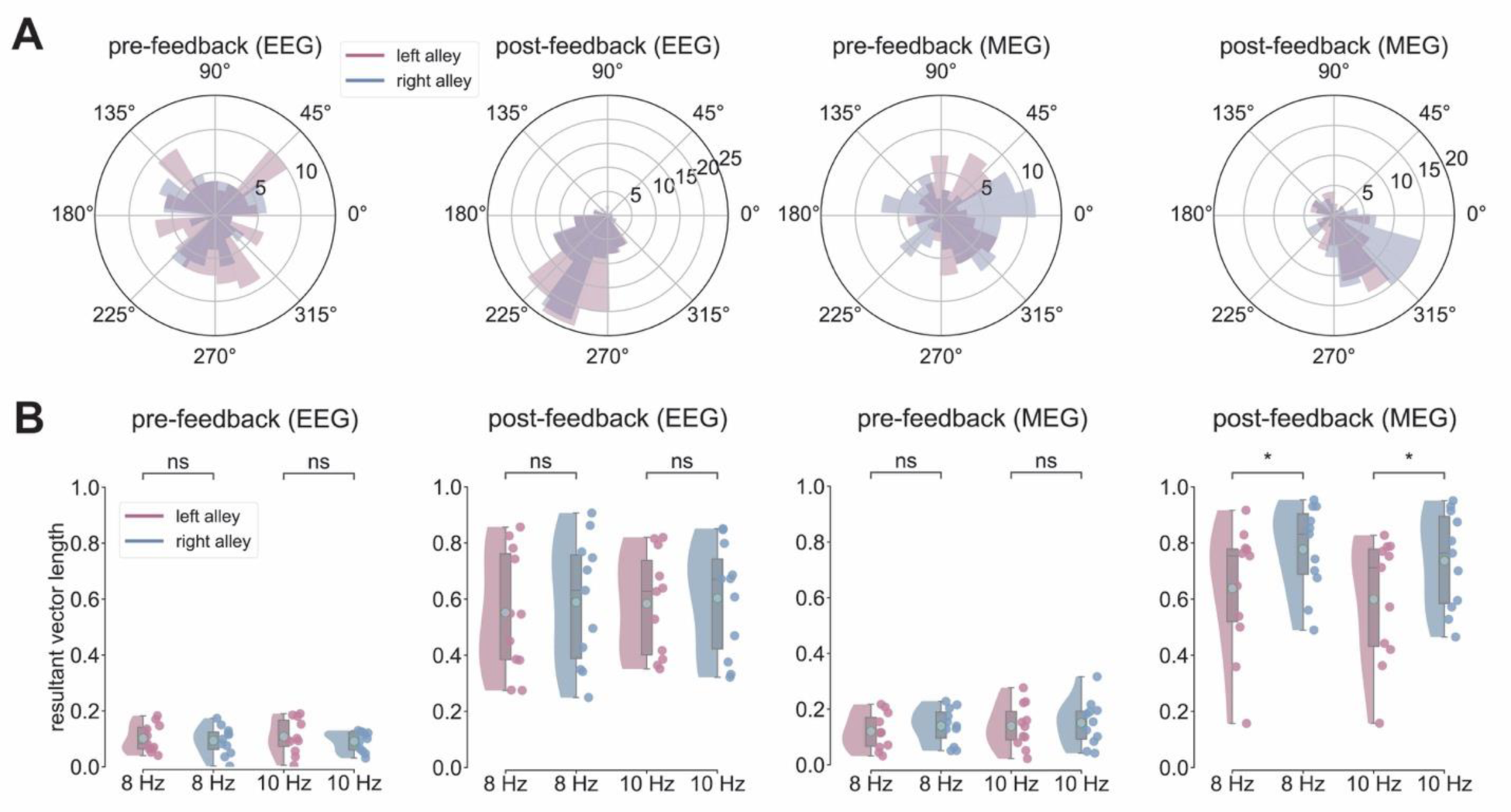
Spatial modulation of phase alignment in EEG and MEG. **A**: Circular distributions of phase angles pre-feedback and post-feedback separated by method (EEG on the left, MEG on the right) and alley of subject 8. Bar size corresponds to the number of trials a particular phase angle was found at the peak timing of the average event-related potential in the EEG and MEG. **B**: Mean resultant vector pre- and post-feedback (EEG left two panels, MEG right two panels) with individual subject values plotted as colored dots separated by alley (left vs. right alley) and frequency (8 Hz vs. 10 Hz). ns: not significant *\*p < 0.05*

When comparing the pre- to post-feedback (±188 ms) histogram for a single example subject’s EEG recording (Figure 4A, left panels), there was a clear alignment of phase angle distributions (see Figure 4B, left panels), with an average CGM aligning at 141° (SD = 35.52°) and an average RVL of 0.58 (SD = 0.19) during the post-feedback window. In the pre-feedback phase alignment, RVL was significantly lower (M = 0.1, SD = 0.03; Figure 4B, left panels) compared to the post-feedback window, *W* = 0, *p* = 0.001, *d* = 3.38. Next, an analysis between left and right alleys in the post-feedback window revealed that right alleys yielded higher RVL values and lower CGM values (8 Hz: MCGM = 180°, SDCGM = 55°, MRVL = 0.59, SDRVL = 0.22; 10 Hz: MCGM = 122°, SDCGM = 72°, MRVL = 0.6, SDRVL = 0.19) than left alleys (8 Hz: MCGM = 160°, SDCGM = 54°, MRVL = 0.55, SDRVL = 0.21; 10 Hz: MCGM = 100°, SDCGM = 77°, MRVL = 0.58, SDRVL = 0.18), but no main effects nor interactions were observed for either measure (p > 0.05).

In regard to MEG recordings, the histogram of phase angles at approximately 196 ms (Figure 4A, right panels) shows a significant alignment of phase angles from pre- (MCGM = 187°, SDCGM = 39°, MRVL = 0.14, SDRVL = 0.05) to post-feedback (MCGM = 173°, SDCGM = 71°, MRVL = 0.69, SDRVL = 0.17), *W* = 0, *p* = 0.003, *d* = 4.17. When comparing left and right alleys in the post-feedback window, the two-way repeated measures ANOVA for RVL data revealed a significant main effect of Feedback Alley Type (*F*(1, 10) = 7.66, *p* = 0.019, *η*^2^= 0.43) such that right alleys displayed the highest concentration of the phase data around the mean direction (8 Hz: MCGM = 163°, SDCGM = 110°, MRVL = 0.78, SDRVL = 0.16; 10 Hz: MCGM = 147°, SDCGM = 113°, MRVL = 0.74, SDRVL = 0.14) compared to left alleys (8 Hz: MCGM = 183°, SDCGM = 116°, MRVL = 0.64, SDRVL = 0.23; 10 Hz: MCGM = 197°, SDCGM = 119°, MRVL = 0.6, SDRVL = 0.22). There was also a significant main effect for Frequency (*F*(1, 10) = 12.51, *p* = 0.005, *η*^2^ = 0.56), with 8 Hz RPT being marked by significantly greater phase alignment than 10 Hz RPT regardless of Feedback Alley Type (left vs. right alley; Figure 4B, right panels). An interaction was not observed (*F*(1, 10) = 0.008, *p* = 0.928, *η*^2^= 0.001), indicating that the difference between right and left alleys was uniform across 8 Hz and 10 Hz. To note, no main effects nor interactions were found when comparing EEG and MEG post-feedback when using CGM as the dependent variable.

## Discussion – Experiment 1

We recently proposed that RPT elicited by reward feedback received during virtual navigation reflects phase resetting of the ongoing theta rhythm in the right PHG, and that the RPT phase shift between feedback in left and right alleys results from differences in the timing of theta phase reset (Baker & Holroyd, 2009, 2013). Here we replicated this RPT effect using both EEG and MEG data recorded from the same group of individuals performing the T-maze task: RPT was stronger and peaked earlier when the feedback-related stimuli were presented in the right alley compared to the left alley. This effect was only observed in sensors over the right hemisphere. Consistent with the partial phase reset hypothesis, ITC analysis in the high theta frequency range (7-10 Hz) yielded a large increase in ITC at the peak timing of RPT in both EEG and MEG datasets, indicating elevated phase consistency across trials. Larger ITC peaks were found for feedback presented in the right alley compared to left alleys in both EEG and MEG datasets, and more so EEG-related ITC values peaked earlier for right alleys compared to left alleys. Moreover, a phase analysis revealed that there was a significant increase in phase alignment at the peak RPT latency compared to baseline for EEG and MEG data, and further, phase alignment was significantly larger for right alleys compared to left alleys in the MEG data, replicating our previous EEG findings (Baker & Holroyd, 2013). Note that we did not replicate the phase alignment difference, as indexed by the RVL, between left and right alleys in the EEG data, possibly due to the small sample size. Together, these EEG and MEG results demonstrate that the RPT occurs as a stimulus-induced increase in power and phase consistency of theta oscillations, the timing (and phase angle) of which depends on the spatial context in which the feedback stimuli are presented (left vs. right alleys).

### Experiment 2: Simultaneous EEG-fMRI

To examine the source of RPT, we recorded EEG-fMRI simultaneously from participants engaged in the Maze/No-maze task used in our previous EEG (Baker & Holroyd, 2013) and fMRI (Baker et al., 2015) experiments (Figure 5). In the “T-maze condition”, participants were asked to find rewards by choosing between two alley options in the virtual maze. In the “No-maze condition”, the participants engaged in a task that was formally identical to the T-maze task except that the imperative and feedback stimuli were displayed against scrambled images of the maze environment, which dissociated these events from their spatial context. This manipulation isolated the contribution of the spatial environment to the RPT effect by holding constant other aspects of task performance such as the degree of motor-related activity. Without a real-life spatial environment like the T-maze, the PHG should not differentiate between feedback following left or right button presses, removing the phase difference coding for spatial location. Hence, we predicted the following: 1) the RPT response would be elicited by the feedback stimuli in both tasks, but the RPT latency effect (earlier RPT peak in right compared to left alley) would occur only in the T-maze condition and not in the No-maze condition (replicating Baker & Holroyd, 2013); 2) feedback stimuli would produce a greater hemodynamic response in the right PHG and other regions associated with virtual navigation (i.e., precuneus and medial temporal cortex) when presented in the T-maze condition compared to the No-maze condition; 3) feedback stimuli presented in the right alley would produce a greater right PHG hemodynamic response compared to the left alley in the T-maze (replicating Baker et al., 2015); and 4) trial-by-trial variation in peak RPT power should be significantly coupled to trial-by-trial variation in PHG BOLD activation, and not to BOLD activation in control regions which are unrelated to spatial navigation or outside of the medial temporal cortex (for further details, see results section “Single-trial right PHG activation is predicted by RPT power” and methods section “Integrated EEG-fMRI analysis”).

**Figure 5.**
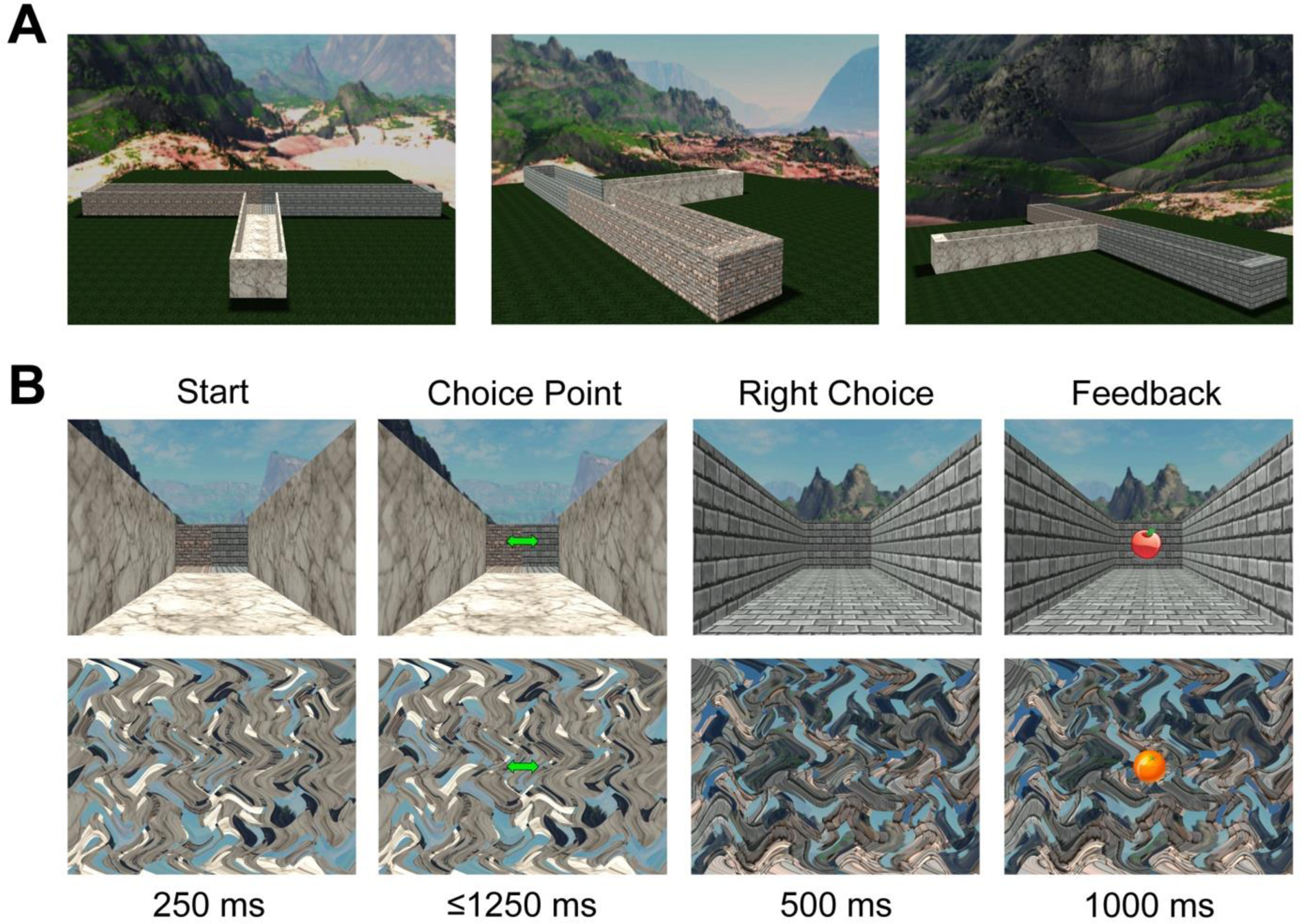
Overview of the Maze/No-maze task. A: Shape of the virtual T-maze as seen from above focused on the start position (left), the left alley (center) and the right alley (right). B: Example trial for Maze-No-maze task used in Experiment 2 (EEG-fMRI) depicting the T-maze (top row) and No-maze condition (bottom row). Headings in the top indicate the stimulus type and the bottom line indicates each stimulus’ duration. Behavioral analyses testing the equivalence of left and right alleys (i.e., reaction times, choice frequency) as well as choice patterns are detailed in Figure 5-figure supplement 1.

### Electrophysiological evidence of the RPT effect in EEG-fMRI

Figure 6 illustrates the results of the time-frequency analysis of the electrophysiological response to feedback encountered in the left and right alley locations. Visual inspection of Figure 6A reveals a clear enhancement of theta power (between 4-12 Hz) peaking approximately 200 ms following the onset of the feedback stimulus in the T-maze (left) and No-maze (right) conditions. This increase in power exhibited a right posterior scalp distribution with a maximum at channels E160 and 107 of the 256-channel EEG geodesic sensor system applied in this experiment, which are equivalents of PO8 and PO7 of the standard 10-20 EEG configuration. These results are characteristic of RPT (e.g., Baker and Holroyd, 2013) and indicate that RPT can be detected when recording EEG during MRI scanning.

**Figure 6.**
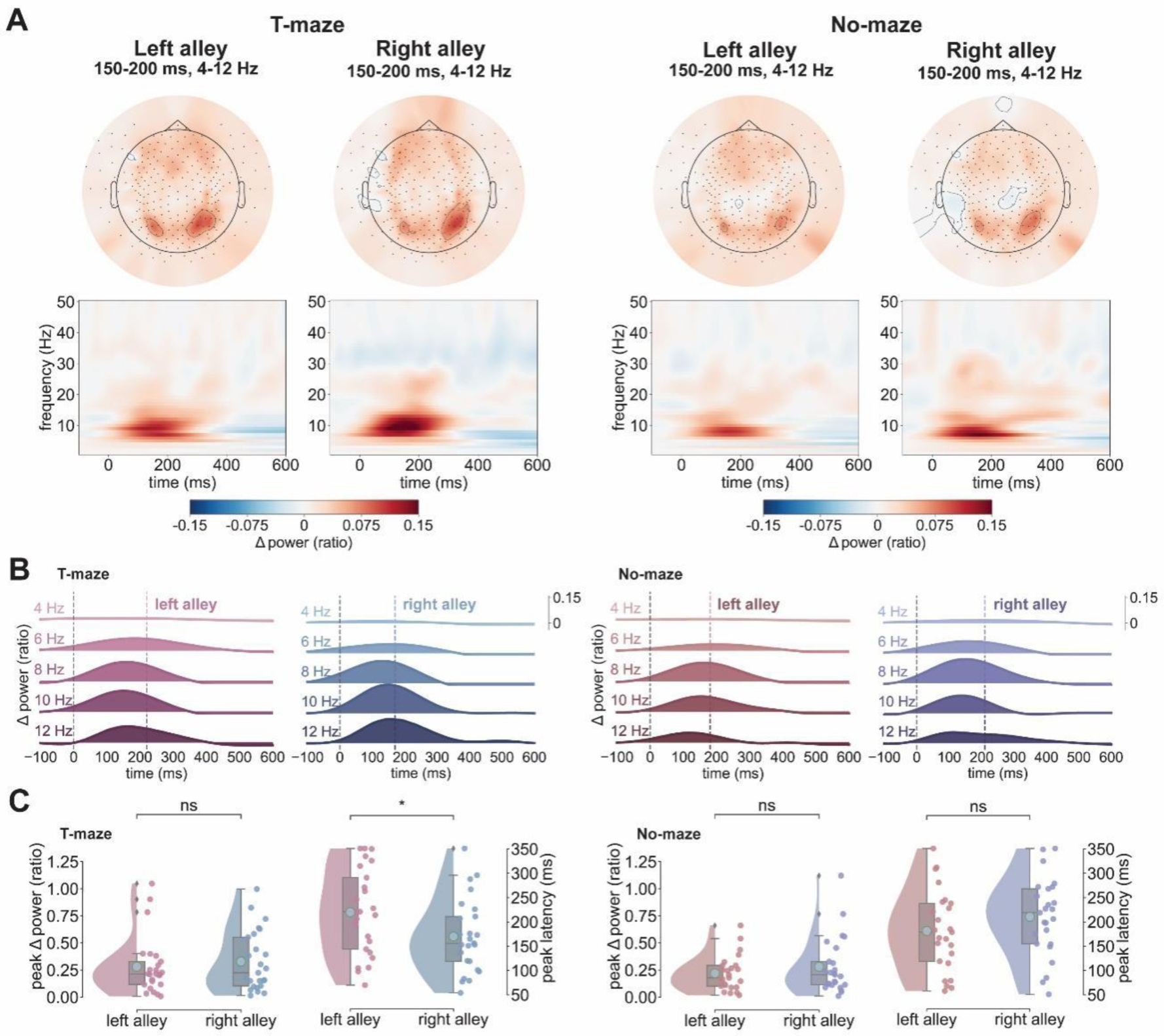
Replication of the RPT effect in simultaneous EEG-fMRI. **A**: Topographies depicting the evoked power for delta (1-4 Hz) and theta bands (4-12 Hz) averaged from 150 ms to 200 ms post-feedback separately for Task Type (T-maze: left panels, No-maze: right panels) and alley (left alleys vs. right alleys). The spectrograms below show the corresponding evoked power from -100 ms to 600 ms at a right parieto-occpital sensor (E160) separated by Task Type and alley. **B**: Ridge plots show the same results as in A divided into the delta band (1-4 Hz) and theta bands (5-6 Hz, 7-8 Hz, 9-10 Hz, and 11-12 Hz), with the average RPT peak timing for each condition marked by a dashed colored line. Results for the T-maze condition are on the left side and for the No-maze condition on the right side. **C**: Raincloud plots depict the RPT power peaks and peak latencies for the T-maze condition on the left and the No-maze condition on the right, with cyan dots showing the mean value of each distribution. Alley is shown on the x-axis and colored dots represent individual subject values. ns: not significant *\*p < 0.05*

In line with our prediction, a repeated measures ANOVA on RPT peak latency as a function of Task (T-maze vs. No-maze) and Feedback Alley Type (left vs. right alley) revealed an interaction between Task and Feedback Alley Type, *F*(1, 24) = 10.14, *p* = 0.003, *η*^2^ = 0.3 (Figure 6B and 6C). A priori comparisons between alleys in the T-maze and No-Maze condition indicated that for the T-maze condition, the RPT peak latency was longer for left alleys (M = 220 ms, SD = 86 ms) compared to right alleys (M = 170 ms, SD = 74 ms), *t*(24) = 2.51, *p* = 0.039 (Bonferroni-Holm corrected), as we predicted. By contrast, in the No-maze condition, no difference was observed for the RPT peak latency to feedback presented following a left button press (M = 181 ms, SD = 81 ms) compared to a right button press (M = 210 ms, SD = 78 ms), *t*(24) = 1.81, *p* = 0.082. Importantly, this T-maze simple main effect was not observed at the contralateral sensor E107 (p > 0.05; see Figure 6-figure supplement 1). A comparable ANOVA on RPT power as a function of Task and Feedback Alley Type revealed a trend for a main effect of Alley Type, *F*(1, 24) = 4.067, *p* = 0.055, *η*^2^ = 0.14, indicating that RPT peak values tended to be larger for right alleys (M = 0.303, SD = 0.263) than for left alleys (M = 0.25, SD = 0.217) regardless of Task Type. No other main effects nor interactions were identified.

### PHG BOLD is modulated by direction only during navigation

Figure 7A depicts the whole brain clusters for the T-maze vs. No-maze contrast from Baker et al. (2015: 1.5T scanner) (left panels) and the current study (right panels: 3.0T scanner) that were significant after False Discovery Rate (FDR) correction. Consistent with our prediction and prior work, feedback stimuli presented in the virtual T-maze task relative to the No-maze task revealed significant activations in bilateral PHG (right: *t*(24) = 6.51, *p* = 0.023, FDR-corrected | left: (*t*(24) = 9.11, *p* = 0.004, FDR-corrected), left temporal-occipital cortex (*t*(24) = 6.53, *p* = 0.026, FDR-corrected), right precuneus (*t*(24) = 5.7, *p* = 0.0001, FDR-corrected) and bilaterally in the cuneus (right: *t*(24) = 8.28, *p* = 0.004, FDR-corrected; left: *t*(24) = 8.83, *p* = 0.026, FDR-corrected), brain regions commonly activated during virtual navigation tasks (Figure 7, Table 1: Epstein, 2008; Kaplan et al., 2014; Ohnishi et al., 2006; Weniger et al., 2010). The No-maze condition displayed a significant cluster in the medial occipital gyrus (*t*(24) = 9.99, *p* = 0.0001, FDR-corrected). A complete overview of whole brain cluster statistics can be found in Table 1. Next, a whole brain analysis between the hemodynamic response to feedback processing in the right alley compared to the left alley of the T-maze revealed activations largely in the motor system (e.g., left primary motor cortex to feedback encountered in the right alley, *t*(24) = 7.4, *p* < 0.0001, FDR-corrected, and activation in the right motor cortex to feedback encountered in the left alley, *t*(24) = 7.36, *p* < 0.001, FDR-corrected), as would be expected given the motor demands of the task and the results of our previous study.

**Figure 7.**
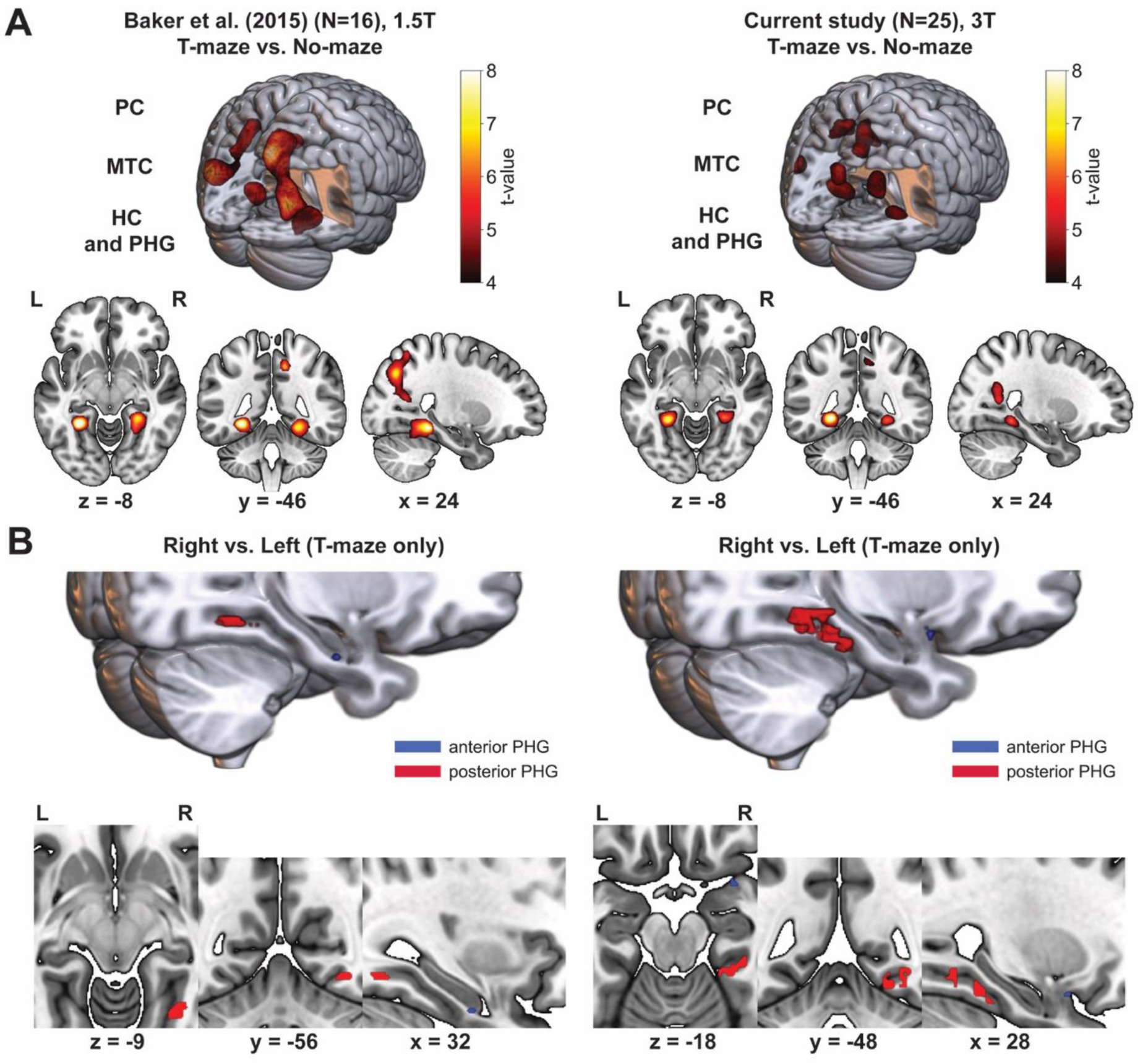
Comparison of the whole brain Maze vs. No-maze contrast and ROI Right vs. Left contrast results between the current and previous analysis. **A**: Left side depicts the data shown in Figure 2B and 2C from Baker et al. 2015 (recorded using a 1.5T scanner). Plots show the results of the whole brain analysis of the maze contrast. The 3D render contains significant clusters stronger during the T-maze condition (p < 0.05, FDR-corrected) in (top right) and outside of the render (top left). Labels indicate the location of the clusters (PC: Precuneus; MTC: Medial Temporal Cortex; PHG: Parahippocampal Gyrus). Multiplanar slices show the color-coded t- values for the same clusters stronger for the T-maze condition in red and yellow (L/R: Left and Right). Analogous plots for the current study depicting the same T-maze vs. No-maze contrast are shown on the right side. **B**: Same as in A, left panels show the ROI PHG results for the T-maze vs. No-maze as well as Right vs. Left (Maze only) contrasts depicted in Figure 3 from Baker et al. 2015. The right side depicts the same contrasts in analogous plots for the results of the current study. 3D renders show the locations clusters specific to the T-maze condition and right alleys specifically in the T-maze condition located in the same anterior (blue; only significant on peak- voxel-level) and the posterior (red; marginally significant on cluster-level) PHG areas across both studies. Multiplanar slices show only the Right vs. Left contrast focused on the bilateral PHG.

**Table 1.**
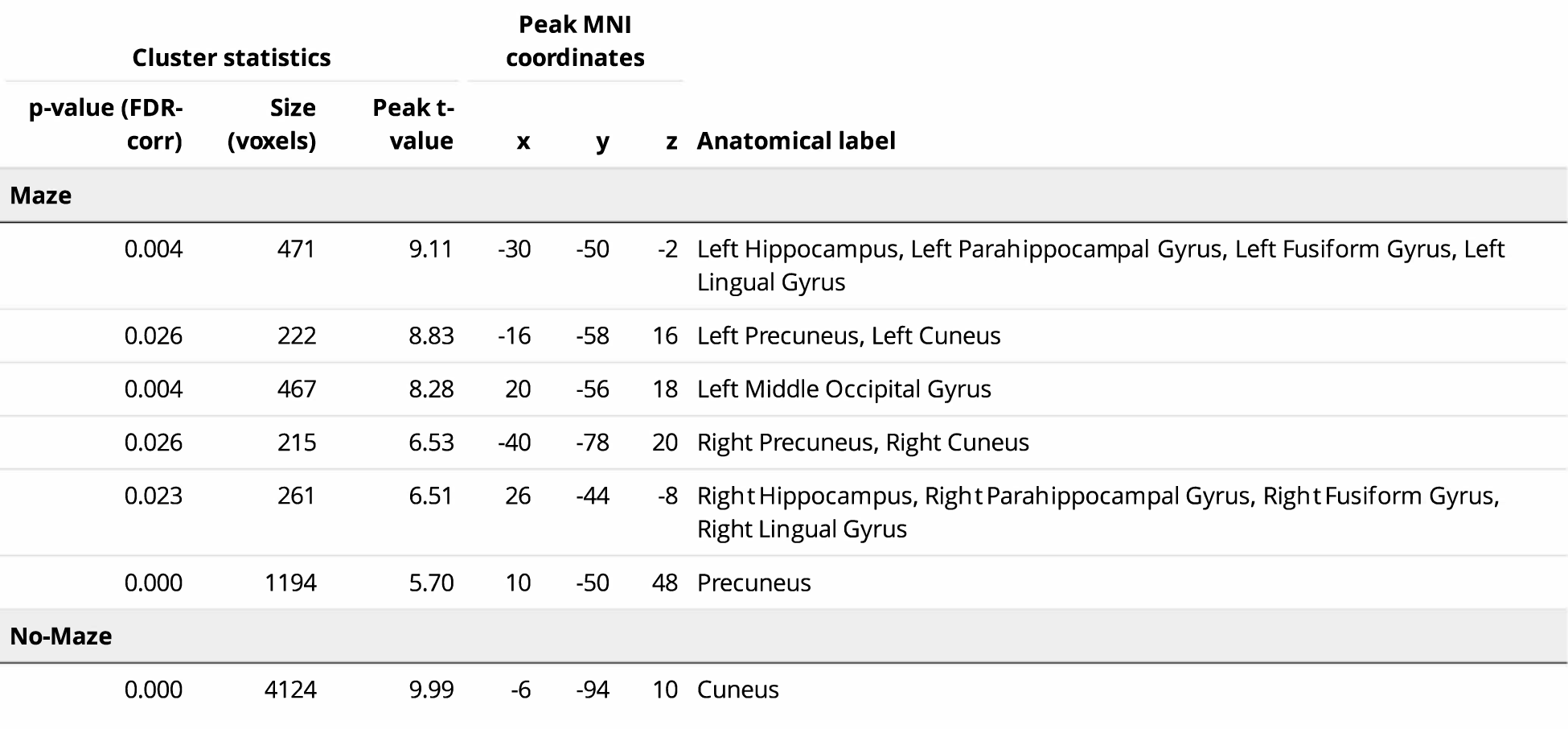
Whole brain cluster statistics for the T-maze vs. No-maze contrast. Rows show significant clusters found to be more strongly activated for the T-maze and No-maze condition respectively.

| Cluster statistics |  |  | Peak MNI coordinates |  |  | Anatomical label |
| --- | --- | --- | --- | --- | --- | --- |
| p-value (FDR-corr) | Size (voxels) | Peak t-value | x | y | z |  |
| Maze |  |  |  |  |  |  |
| 0.004 | 471 | 9.11 | -30 | -50 | -2 | Left Hippocampus, Left Parahippocampal Gyrus, Left Fusiform Gyrus, Left Lingual Gyrus |
| 0.026 | 222 | 8.83 | -16 | -58 | 16 | Left Precuneus, Left Cuneus |
| 0.004 | 467 | 8.28 | 20 | -56 | 18 | Left Middle Occipital Gyrus |
| 0.026 | 215 | 6.53 | -40 | -78 | 20 | Right Precuneus, Right Cuneus |
| 0.023 | 261 | 6.51 | 26 | -44 | -8 | Right Hippocampus, Right Parahippocampal Gyrus, Right Fusiform Gyrus, Right Lingual Gyrus |
| 0.000 | 1194 | 5.70 | 10 | -50 | 48 | Precuneus |
| No-Maze |  |  |  |  |  |  |
| 0.000 | 4124 | 9.99 | -6 | -94 | 10 | Cuneus |

As before, we conducted a ROI analysis focusing exclusively on the right and left PHG (Figure 7B; left panel: results from Baker et al. 2015, right panel: current results). Consistent with our previous study, bilateral clusters were observed in a posterior region of the right PHG (*t*(24) = 6.51, *p* = 0.026, FDR-corrected) and the left PHG (*t*(24) = 8.67, *p* = 0.023, FDR-corrected) when contrasting feedback processing in the T-maze condition versus the No-maze condition. The same posterior cluster (*t*(24) = 4.4, *ppeak-level* = 0.0001, uncorrected, *pcluster-level* = 0.049, uncorrected) and an anterior cluster (*t*(24) = 3.2, *ppeak-level* = 0.002, uncorrected, *pcluster-level* = 0.75, uncorrected) emerged in the right hemisphere when subjects chose the right alley as opposed to left alley in the T-maze. The opposite contrast revealed no voxel values that were stronger for feedback stimuli presented in the left alley compared to the right alley of the maze, and an analysis on the left PHG ROI did not reveal any significant activation for either the left alley-right alley or right alley-left alley contrasts. These clusters directly mirrored the results from Baker et al. (2015) as can be seen when comparing the left and right panels of Figure 7B, highlighting the PHG’s robust engagement during feedback processing in a spatial environment.

### Single-trial right PHG activation is predicted by RPT power

Building on these unimodal results, the goal of the integrated EEG-fMRI analysis was to take advantage of the complementary temporal and spatial resolution of each method. By integrating the EEG and fMRI data, we hoped to determine whether or not RPT reflected a PHG theta reset mechanism for spatial navigation. It is important to note that the PHG encompasses a large portion of the medial temporal lobe, and that it has been traditionally accepted to include different regions along its anterior-posterior axis (i.e., entorhinal cortex, perirhinal cortex, and parahippocampal cortex). In addition, a significant body of research has identified several functional processes linked to the parahippocampal region (Aminoff et al., 2013; Epstein, 2008; Maguire, Frith, et al., 1998; Owen et al., 1996; Weniger & Irle, 2006). Furthermore, given the highly functional and interconnected nature of the parahippocampal region, it has recently been argued that its anatomically defined subregions should be further subdivided by its connectivity with other cortical and subcortical areas (Syversen et al., 2021). This was motivated by the conflicting results stemming from cross-species comparisons between humans, rodents and non-human primates, as well as from human neuroimaging data on spatial memory processing, and from observations of neurodegenerative structural changes of human PHG divisions based on animal cytoarchitecture and landmarks (Syversen et al., 2021). It is likely that such conflicting results arise from variable definitions of parahippocampal sub-regions across studies. We therefore proposed that a well-defined structural connectivity-based segmentation of the human PHG was a necessary precursor for understanding how spatial contextual information is processed by the PHG during goal-directed navigation, and further, provide a more precise localization of RPT activity (Baker, Reid, et al., 2017). Thus, redefining the conventional PHG ROI using formulating criteria based both on anatomical landmarks and connectivity profiles was relevant for the interpretation of the current EEG-fMRI integration analysis, as well as for future studies examining the functional relevance of its subregions in goal-directed behavior.

Here, we used probabilistic tractography data from the Human Connectome Project (HCP) to segment the anterior and posterior PHG based on differential connectivity profiles to the whole brain region (Baker, Reid, et al., 2017). A detailed description of the tractography-based PHG parcellation analysis and results can be found in the supplementary section “PHG segmentation analysis: Diffusion-weighted and resting-state functional MRI”. In brief, the boundaries of the anterior PHG (aPHG) and posterior PHG (pPHG) region were first defined by reference to the morphology of the sulci of the collateral sulcal complex as described by Huntgeburth and Petrides (2012) (Figure 8A). Once defined, the aPHG and pPHG masks were linearly registered to native diffusion space and we then applied a data-driven connectivity-based brain parcellation procedure (i.e., spectral clustering) described in Zhang et al. (2017) and Fan et al. (2016). This method delineates the boundary of subdivisions based on the whole brain connectivity fingerprints of each voxel in the seed region (here the anterior and posterior PHG; Figure 8). The tractography-based parcellation approach yielded 4 subregions within the right aPHG, and 2 subregions within the right pPHG (Figure 8A and 8B). A similar number of subregions were also obtained for the left aPHG and pPHG, demonstrating a symmetric pattern between left and right hemispheres. These PHG clusters exhibited robust cortico-cortical connections (e.g., frontal, occipital, parietal, and temporal) and cortico-subcortical connections (e.g., hippocampus and basal ganglia), demonstrating a caudal-to-rostral trend of fiber projections. Furthermore, the connectivity patterns for each subregion were highly consistent with resting-state functional connectivity (RSFC) patterns obtained from the same neuroimaging data (Figure 8C and 8D).

**Figure 8.**
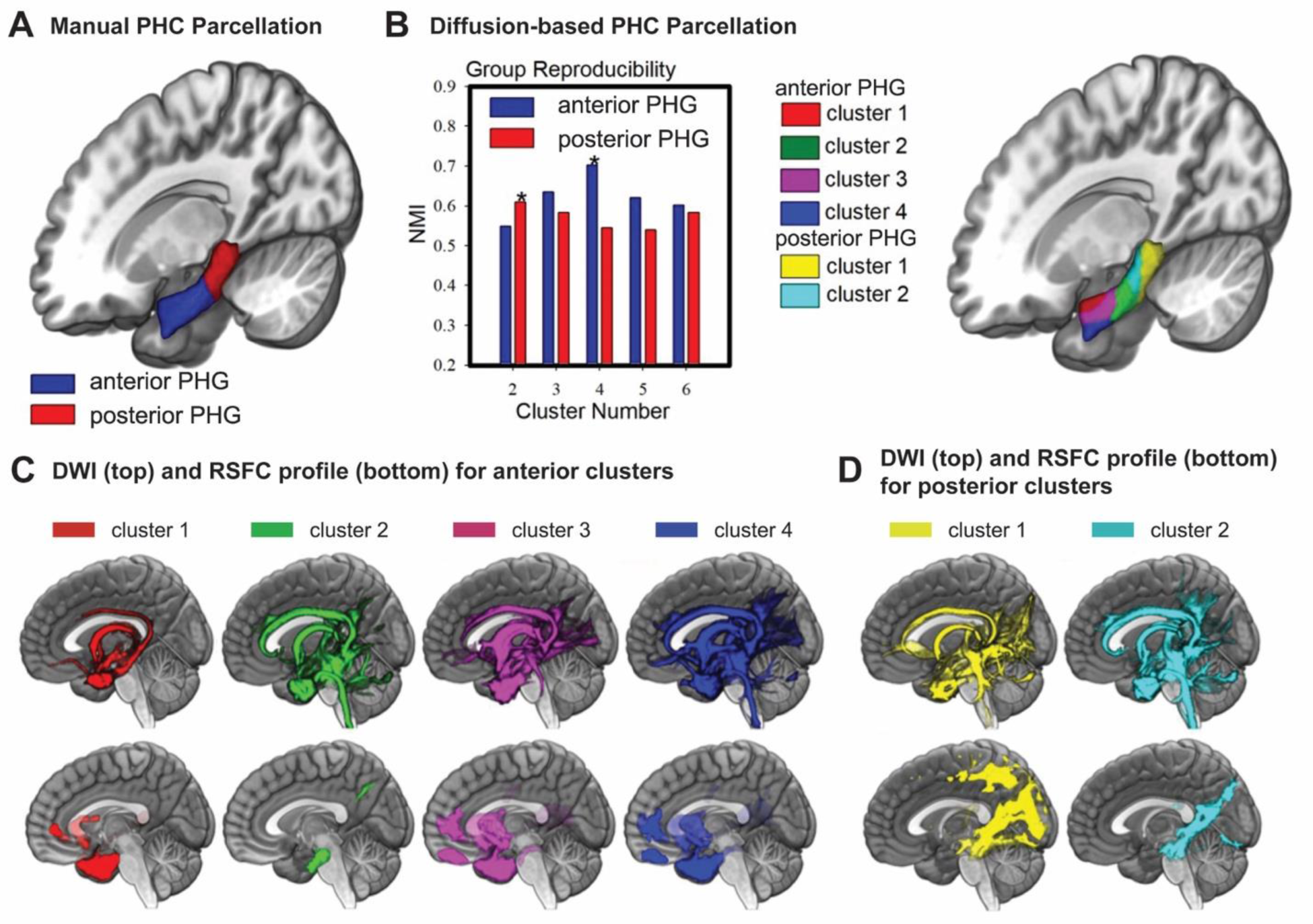
Structural and functional connectivity profile of the PHG. **A**: Illustration of the PHG manual parcellation map of the anterior (in blue) and posterior (in red) PHG as established in the present article. **B**: Anterior and posterior PHG diffusion parcellation results for the HCP datasets. X-axis shows size of cluster solution (i.e., two- to six-cluster solution). Normalized mutual information provides an indication of clustering consistency in which high values indicate good consistency (left panel). The figure shows that the four-cluster solution had the highest normalized mutual information value for the anterior PHG, and two-cluster solution for the posterior PHG (marked by stars). Surface view of the PHG parcellation results (right panel). **C**: Population maps of the probabilistic tractography (top row) and RSFC (bottom row) results for each right anterior cluster. Statistical parametric maps are displayed using a statistic thresholded at p < 0.001 (FDR-corrected) with the intensity scales representing a T-value of 18. All figure maps are shown on the ICBM152 template in MNI space with MRIcroGL. **D:** Population maps analogous to **C** for the posterior clusters.

Next, the integrated EEG-fMRI analysis focused on single-trial beta coefficients from the bilateral PHG ROIs and the averaged evoked power at channel E160 during T-maze and No-maze trials. When averaging single-trial power estimates for the T-maze and No-maze condition across left and right alleys, these electrodes carried the most pronounced RPT power in their respective hemisphere. To test the link between the EEG and fMRI data, a set of multimodal Linear Mixed Effects (LME) models were estimated with single-trial BOLD averages for each of the twelve bilateral PHG ROIs as the dependent variables. Thus, one model was run for each ROI in each condition (T-maze vs. No-maze). Five centered single-trial EEG power estimates (mean power from 50 ms to 250 ms post-feedback) were used to create delta (1-4 Hz) and theta (4-12 Hz) band regressors for each subject that were included in each model. Resulting fixed effects estimates were standardized to beta weights and tested for significance. To note, because prevailing definitions of the human (4-8 Hz) (Ekstrom et al., 2005) and animal (4-12 Hz) (O’Keefe, 1993; O’Keefe & Recce, 1993) theta rhythms are inconsistent with one another, and because our previous time-frequency results showed a prominent upper theta range (7-10 Hz), we decided to split the theta range into four regressors with a bandwidth of 2 Hz each (5-6 Hz, 7-8 Hz, 9-10 Hz, and 11- 12 Hz). All p-values were corrected for multiple comparisons using the FDR method for each regressor within a model.

To test directly our primary hypothesis that single-trial RPT power can predict single-trial PHG BOLD values, we specifically tested the relationship specifically between RPT regressors and BOLD signal from the six PHG ROIs on the right hemisphere (full LME model results for both hemispheres can be found in Table 2-table supplements 1-5). As a control and to differentiate against slower oscillations involved in feedback processing, we also included a delta power regressor in each model. Single-trial BOLD values in aPHG2 (*t* = 2.81, *p* = 0.03, *b* = 0.129), pPHG1 (*t* = 2.691, *p* = 0.043, *b* = 0.123), and pPHG2 (*t* = 3.207, *p* = 0.008, *b* = 0.148) were significantly predicted by 7-8 Hz evoked power recorded from the right parietooccipital EEG sensor E160 (Figure 9). The 7-8 Hz regressor received a positive weight for both ROIs, indicating that for a given trial increases in PHG activation were coupled to increases in 7-8 Hz power. The frequency band below, 5-6 Hz, was also significantly coupled to aPHG2 (*t* = -2.559, *p* = 0.032, *b* = -0.097), and pPHG2 (*t* = -2.522, *p* = 0.024, *b* = -0.097) but with each regressor receiving a negative weight. Hence, a trial-to-trial increase in activation across these PHG ROIs was associated with a decrease in 5-6 Hz power. An overview of all beta weights for the 5-6 Hz and 7- 8 Hz theta bands plotted onto their respective ROIs can be found in Figure 9-figure supplement 1. None of the remaining delta and theta bands received significant beta weights. Table 2 summarizes the full results for the right PHG models (left PHG models can be found in Table 2-table supplement 1). Importantly, among the No-maze condition LME models, none of the aforementioned relationships between single-trial RPT power and PHG BOLD activation were found (see Table 2-table supplement 2 and Table 2-table supplement 3), suggesting that the RPT- PHG associations predominantly emerged during spatial (vs. non-spatial) feedback. As a control analysis, we used the same RTP power regressors to predict single-trial BOLD activation in the precuneus (whole brain cluster from Figure 7A) and the middle superior temporal gyrus (i.e., Heschl’s Gyrus). Neither exhibited significant relationships with RPT power regressors (Table 2- table supplement 4), demonstrating that the above couplings were specific to the PHG ROIs.

**Figure 9.**
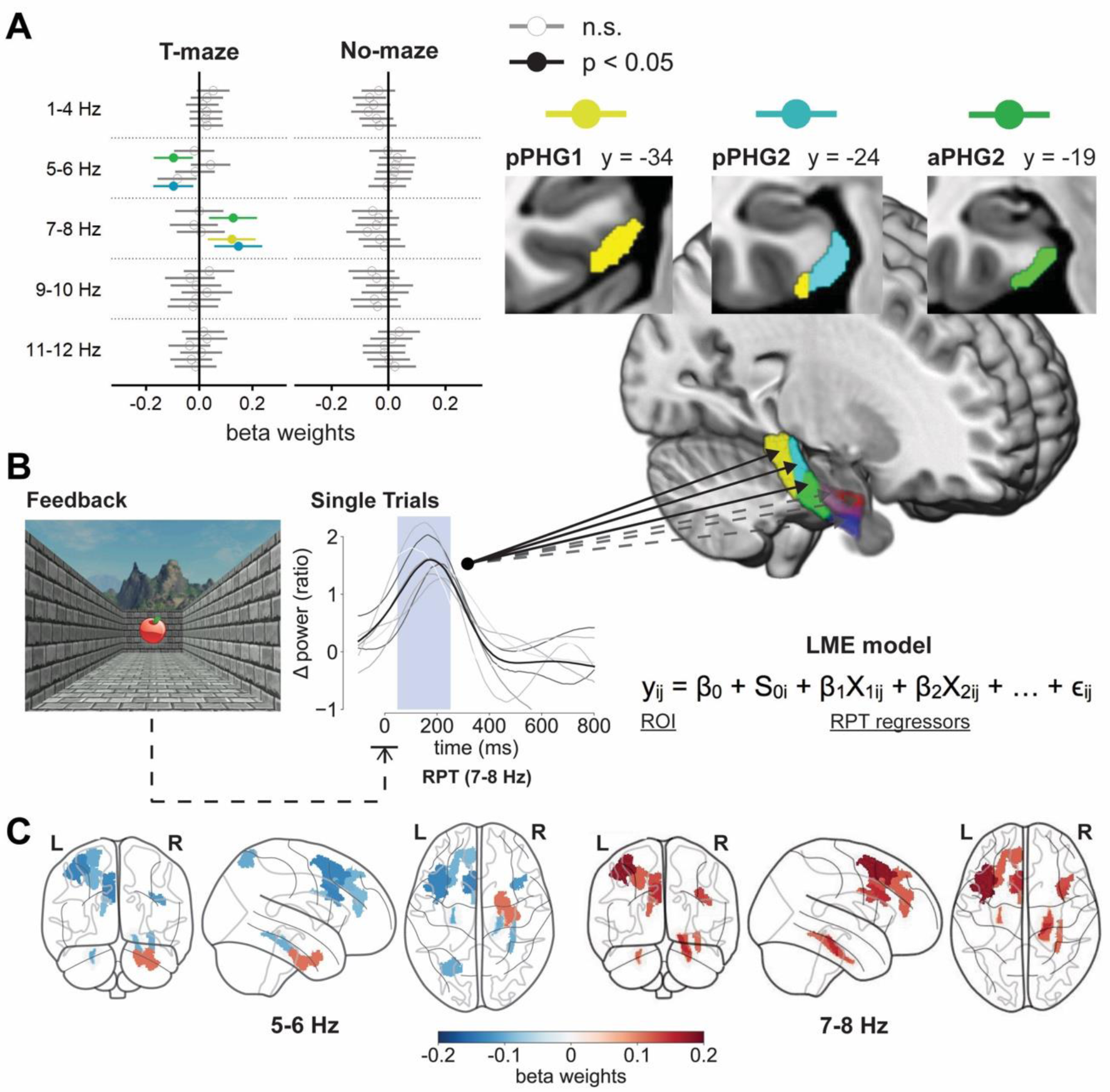
Results from multimodal LME models predicting single-trial BOLD using RPT regressors. A: Forest plots on the left depict the beta weights for the EEG-based regressors by frequency band and PHG ROI (aPHG2: green, pPHG1: gold, pPHG2: light blue). Left column shows the results for the T-maze condition, right column shows the results for the No-Maze condition. The position of circles on the x-axis on each forest plot reflects the size of each weight. Filled out and colored circles are significant beta weights with a p-value < 0.05 after FDR correction. Gray, empty beta weights are not significant. Lines around dots represent a 95% confidence interval for the respective weight. Sagittal view and coronal slices on the right show the PHG clusters that were significantly predicted by at least one theta frequency band. **B:** Bottom panels represent a schematic overview and the equation of the LME models. Line plots depict feedback-locked example trials recorded at E160 (gray, transparent lines) and their average (black line) for subject 8. The mean amplitude from 50 ms to 250 ms (blue window) was used to build regressors. In each model, y (BOLD value for a given subject i and trial j) was predicted by the overall model intercept β0, a random intercept S0 for each subject, a fixed regressor β1-5 for each EEG frequency band that is weighted by the power for a given trial j and subject i, and a corresponding error term ɛ. Solid arrows denote significant relationships between single-trial EEG regressors and BOLD signals, dashed and transparent arrows denote non-significant relationships. **C:** Color-coded PHG beta weights from the LME models run for each Glasser ROI along with the PHG ROIs. Locations of PHG ROIs created in this study (Figure 8B) are highlighted and encircled in gray. Results are separated by weights for the 5-6 Hz regressors (left) and 7-8 Hz regressors (right). All weights are for the T-maze condition.

**Table 2.**
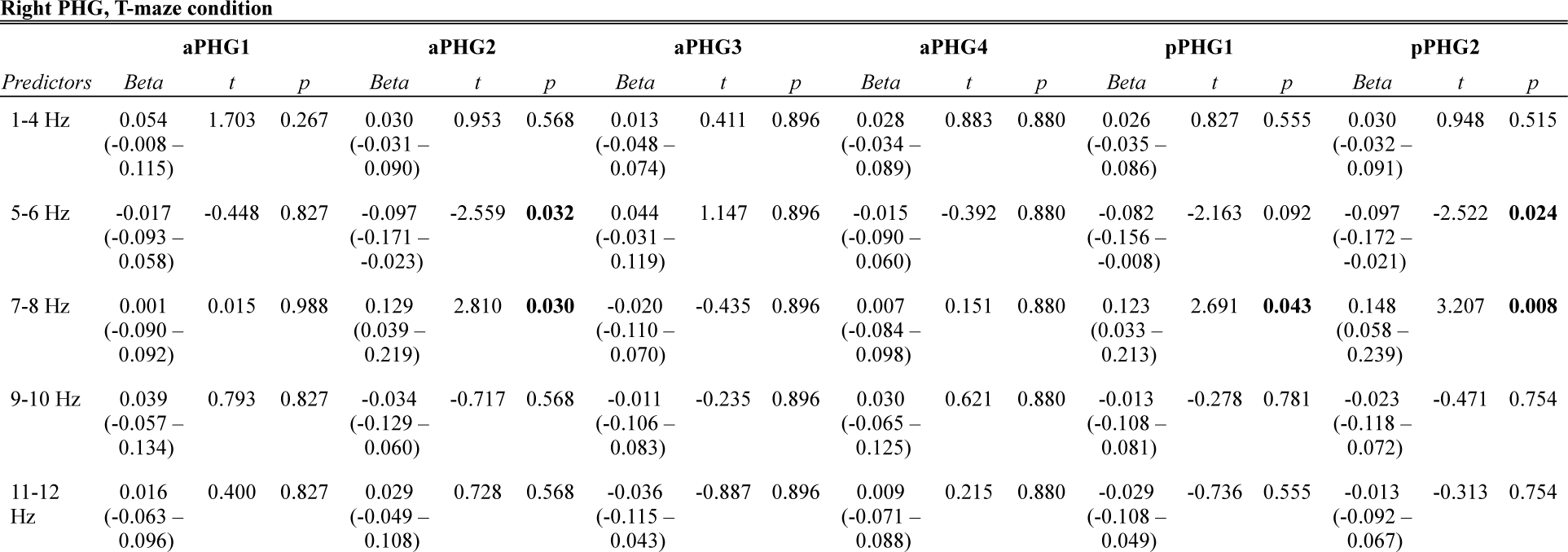
Fixed Effects table of the LME model run on single-trial right PHG ROI BOLD activation (columns) recorded during the T-maze condition. Fixed Effect predictors include four EEG regressors (rows) based on evoked delta power (1-4 Hz) and four RPT bands (5-6 Hz, 7-8 Hz, 9- 10 Hz, 11-12 Hz). All listed p-values were corrected for multiple comparisons within a model. Significant p-values (p < 0.05, FDR-corrected) are printed bold next to the corresponding t-value. For each weight a 95% Confidence Interval (CI) is given below the standardized beta estimate.

| Predictors | aPHG1 |  |  | aPHG2 |  |  | aPHG3 |  |  | aPHG4 |  |  | pPHG1 |  |  | pPHG2 |  |  |
| --- | --- | --- | --- | --- | --- | --- | --- | --- | --- | --- | --- | --- | --- | --- | --- | --- | --- | --- |
|  | Beta | t | p | Beta | t | p | Beta | t | p | Beta | t | p | Beta | t | p | Beta | t | p |
| 1-4 Hz | 0.054<br>(-0.008 – 0.115) | 1.703 | 0.267 | 0.030<br>(-0.031 – 0.090) | 0.953 | 0.568 | 0.013<br>(-0.048 – 0.074) | 0.411 | 0.896 | 0.028<br>(-0.034 – 0.089) | 0.883 | 0.880 | 0.026<br>(-0.035 – 0.086) | 0.827 | 0.555 | 0.030<br>(-0.032 – 0.091) | 0.948 | 0.515 |
| 5-6 Hz | -0.017<br>(-0.093 – 0.058) | -0.448 | 0.827 | -0.097<br>(-0.171 – -0.023) | -2.559 | **0.032** | 0.044<br>(-0.031 – 0.119) | 1.147 | 0.896 | -0.015<br>(-0.090 – 0.060) | -0.392 | 0.880 | -0.082<br>(-0.156 – -0.008) | -2.163 | 0.092 | -0.097<br>(-0.172 – -0.021) | -2.522 | **0.024** |
| 7-8 Hz | 0.001<br>(-0.090 – 0.092) | 0.015 | 0.988 | 0.129<br>(0.039 – 0.219) | 2.810 | **0.030** | -0.020<br>(-0.110 – 0.070) | -0.435 | 0.896 | 0.007<br>(-0.084 – 0.098) | 0.151 | 0.880 | 0.123<br>(0.033 – 0.213) | 2.691 | **0.043** | 0.148<br>(0.058 – 0.239) | 3.207 | **0.008** |
| 9-10 Hz | 0.039<br>(-0.057 – 0.134) | 0.793 | 0.827 | -0.034<br>(-0.129 – 0.060) | -0.717 | 0.568 | -0.011<br>(-0.106 – 0.083) | -0.235 | 0.896 | 0.030<br>(-0.065 – 0.125) | 0.621 | 0.880 | -0.013<br>(-0.108 – 0.081) | -0.278 | 0.781 | -0.023<br>(-0.118 – 0.072) | -0.471 | 0.754 |
| 11-12 Hz | 0.016<br>(-0.063 – 0.096) | 0.400 | 0.827 | 0.029<br>(-0.049 – 0.108) | 0.728 | 0.568 | -0.036<br>(-0.115 – 0.043) | -0.887 | 0.896 | 0.009<br>(-0.071 – 0.088) | 0.215 | 0.880 | -0.029<br>(-0.108 – 0.049) | -0.736 | 0.555 | -0.013<br>(-0.092 – 0.067) | -0.313 | 0.754 |

Finally, to investigate the specificity of this RPT and PHG coupling, we extended the analysis to the whole brain level by running LME models with the same four EEG regressors predicting single-trial BOLD activation in the T-maze condition for each ROI in the Glasser atlas (Glasser et al., 2016). If RPT power is specifically connected to the PHG and spatial navigation, EEG-based RPT regressors should not predict any nearby temporal or parietal regions unrelated to spatial navigation (e.g., superior temporal gyrus). Figure 9C highlights the significant results of this whole brain analysis for all Glasser ROIs in addition to our PHG ROIs specifically for the two previously significant theta bands, 5-6 Hz and 7-8 Hz (for all regressors, see Table 2-table supplement 5 and Table 2-table supplement 6). Almost all signs matched the PHG results, with all but one 5-6 Hz weights receiving a negative sign and all 7-8 Hz weights receiving positive signs. In the right hemisphere two memory-related structures in the medial temporal lobe laterally to the the PHG had significant beta weights: one laterally adjacent to our pPHG definition (Glasser label: “Parahippocampal Area 3”) for both 5-6 Hz (*t* = -2.453, *p* = 0. 045, *b* = -0.097) and 7-8 Hz (*t* = 2.435, *p* = 0. 045, *b* = 0.112) and one in the perihinal cortex (“Ectorhinal-Perhinal Cortex”) only for 5-6 Hz (*t* = 2.898, *p* = 0. 023, *b* = 0.109). Note that the Glasser atlas employs an anatomical delineation of the PHG that laterally extends into and past the collateral sulcus and thus includes regions that we did not include in our definition (e.g., “Parahippocampal Area 3”). No other significant relationships were identified in the temporal and parietal lobe for the right hemisphere. Furthermore, six left hemispheric and two right prefrontal regions, including areas relevant to feedback processing such as the left anterior cingulate cortex and the dorsolateral prefrontal cortex, displayed significant relationships with 5-6 Hz and 7-8 Hz (for full results, see Table 2-table supplement 5 and Table 2-table supplement 6).

## Discussion – Experiment 2

We recently demonstrated that the presentation of feedback-related stimuli in a virtual environment induces a burst of EEG oscillations in the theta frequency range over right posterior areas of the human scalp (RPT), and that the power and phase angle of RPT was significantly larger when participants receive feedback following right turns in the maze compared to left turns in the maze. Even though source localization procedures have indicated that the RPT is produced in or near the PHG (Baker & Holroyd, 2013), and, even though the tasks previously employed to produce the RPT effect have generated a strong hemodynamic response in the right PHG (Baker et al., 2015), the origin of RPT still remains uncertain. This may be due to the inherent spatial limitations of EEG recordings and the lack of time-frequency information in fMRI data. These limitations are formally insoluble but can be ameliorated by the application of the “converging methods” approach in which multiple source analysis techniques are utilized to compensate for their respective weaknesses (Luck, 1999, 2005, p. 296). We hypothesized that converging evidence using simultaneous EEG-fMRI data recorded during the virtual T-maze task would provide strong evidence that RPT is in fact generated in the PHG.

First, to test whether RPT depends on spatial processing rather than other aspects of task performance, we had participants engage in two tasks: the standard T-maze task and a No-maze task. The No-maze task was formally identical to the T-maze but was devoid of the spatial context. As predicted, the RPT effect (latency difference between left and right alleys) occurred only when participants processed the feedback in the maze environment, ruling out other potential sources of the effect such as dependence on right vs. left button presses. The power of RPT was equivalent across the two tasks, suggesting that the underlying neural system was fully engaged even in the No-maze task. Thus, despite the small size of the latency difference (approx. 50 ms), these results replicate our previous work (Baker & Holroyd, 2009, 2013). Notably, several studies have shown that neighboring regions of the PHG, particularly the fusiform gyrus (Iidaka et al., 2006; Rossion et al., 2003), play an important role in object recognition. We proposed that RPT was generated by this network for object processing in the ventral-medial temporal cortex, and that the associated right-hemispheric dominance and latency effect occurred when this processing stream was modulated by spatial context.

Second, consistent with our predictions for the fMRI results we found that the right PHG was more strongly activated by feedback stimuli encountered in the same virtual T-maze that we utilized in our previous EEG studies, when compared to feedback stimuli presented with no spatial context (i.e., No-maze task). Further, statistically significant activations were also observed in the left PHG, bilaterally in the middle temporal-occipital cortex, and in the right precuneus (Figure 7, Table 1), regions commonly activated during virtual navigation, and consistent with our previous report (Baker et al., 2015). In particular, precuneus activity in fMRI studies has been associated with spatial navigation (Lithfous et al., 2017; Ohnishi et al., 2006), self-referential representations during episodic memory (Hebscher et al., 2018), mental imagery (Cavanna & Trimble, 2006) and it has been reported as a part of the default-mode network (Utevsky et al., 2014). It is often associated with allocentric spatial memory and memory retrieval (Frings et al., 2006; Wallentin et al., 2008; H. Zhang & Ekstrom, 2013), but also with the updating of egocentric spatial information that pertains to an individual’s position relative to motivationally relevant objects (Dordevic et al., 2022; Galati et al., 2000).

Third, we predicted that in the T-maze the right PHG would generate a stronger hemodynamic response to feedback stimuli encountered in the right alley compared to the left alley of the T-maze. Our results confirmed this prediction. Consistent with our previous fMRI study, the current analysis revealed two distinct regions within the right PHG: in the T-maze versus No- maze contrast, a large significant cluster was observed in the pPHG, whereas in the right alley vs. left alley contrast, a large pPHG and a small aPHG cluster of activation were noted. In contrast to the viewpoint that the PHG is a single coterminous region that performs a unitary spatial function, our findings support the idea that the PHG is composed of multiple functional components along its posterior-anterior axis. For example, the Parahippocampal Place Area (a region reported to occupy the posterior portion of the PHG and extending into the lingual gyrus) responds preferentially to pictures of places (Epstein et al., 1999, 2003), the posterior and middle regions of the PHG are involved in the encoding and retrieval of information relevant to navigation (Huntgeburth et al., 2017; Janzen & Van Turennout, 2004; Sulpizio et al., 2013), and the entorhinal cortex (occupying the anterior portion of the PHG) has been shown to play a more general role in memory (Mankin et al., 2021; Owen et al., 1996). Moreover, evidence from functional connectivity studies supports the idea that aPHG and pPHG exhibit distinct connectivity patterns, with the pPHG exhibiting a stronger relationship with visual regions in the occipital lobe and the aPHG being more strongly coupled with parietal lobe regions implicated in spatial processing (Baldassano et al., 2013; D. M. Watson & Andrews, 2024; Zhuo et al., 2016). Consistent with this view, our analysis of an HCP dataset showed that subregions within the PHG were diversly connected to unimodal and polymodal association areas as well as frontal regions. In particular, this analysis revealed connections that included the medial frontal cortex and anterior midcingulate via the cingulum bundle, as well as the medial occipital and temporal cortices and the precuneus via the posterior corona radiata.

The above detailed analyses served largely to replicate our previous EEG and fMRI findings. In addition, the main purpose of the study was to test whether trial-by-trial variation in RPT power was coupled to trial-by-trial variation in PHG BOLD activation, which would implicate the right PHG in producing RPT. Consistent with our prediction, we observed that trial-by-trial variation in theta power in the 7-8 Hz range predicted trial-by-trial fMRI activation in two clusters in the right pPHG and one cluster in the right aPHG. This result suggests that with increased BOLD signals in the PHG, subjects exhibited enhanced power in RPT (7-8 Hz), supporting the hypothesis that the identified PHG clusters are the plausible generators of RPT recorded at the scalp. Importantly, there were no significant relationships between RPT and PHG activation during the No-maze condition in any of the LME models. This highlights that the theta-linked PHG engagement may be specific to spatial navigation. Furthermore, in our control and whole brain analyses, most significant couplings between BOLD signal and RPT in the right temporal and parietal cortical areas were exclusive to the medial temporal lobe, demonstrating that the significant RPT and BOLD couplings were specific to our hypothesized regions. Significant relationships between RPT and primarily left prefrontal regions (e.g., anterior cingulate cortex, dorsolateral prefrontal cortex) likely stemmed from these areas being co-activated with the PHG due to their involvement in feedback processing (Baker, Lesperance, et al., 2017; Becker et al., 2014; X. Liu et al., 2011; Marco-Pallarés et al., 2007; Sescousse et al., 2013). However, given their distance to the electrode site where RPT is recorded (E160), these cortical areas are unlikely sources of RPT. Instead, they could be part of the spatial navigation network activated as a result of phase resetting in the PHG propagating spatial feedback information. This idea is supported by our findings of strong structural and functional connectivity between our PHG ROIs and prefrontal cortices.

An unexpected finding in this study was that RPT did not yield a uniform significant relationship with PHG BOLD across individual RPT bands. 5-6 Hz was significantly negatively coupled to almost the same ROIs that were positively associated with 7-8 Hz, suggesting that the unique variances of 5-6 Hz and 7-8 Hz were inversely related to PHG BOLD, where 7-8 Hz increased with heightened BOLD on a given trial while 5-6 Hz decreased. A possible explanation is that phase resetting could result in the synchronization of oscillations in the preferred frequency range of 7-8 Hz for optimal encoding of reward locations, while the immediately adjacent frequencies like 5-6 Hz on a trial-by-trial level are suppressed. The need for the tuning of encoding-related theta frequencies was recently demonstrated by Quirk et al. (2021) who created memory encoding deficits in mice searching a maze for navigational cues by shifting the frequency of endogenous theta rhythm of various hippocampal and entorhinal cell types, such as grid cells. Pushing the endogenous theta rhythms of grid cells (6-9 Hz) above their preferred range decreased the animals’ ability to encode reward information. Such frequency tuning towards an optimal range has also been reported for cognitive control in humans (Senoussi et al., 2022). In addition, the 5-6 Hz RPT power as observed in our experiments could stem at least partially from entorhinal and perirhinal cortex activity, which in our whole brain analysis was the only ROI with a positive coupling to 5-6 Hz, while the PHG’s rhythm gravitated towards 7-8 Hz at the time of the phase reset.

This evidence along with the current whole brain fMRI results and the coupling of RPT power to ROIs (aPHG2, pPHG1, pPHG2) with strong connectivity with the precuneus suggests that phase resetting could be occurring specifically in subregions of the PHG processing egocentric spatial information related to rewards. These regions might exchange information with the precuneus to embed egocentric feedback information into larger spatial contexts and update the individual’s position relative to the reward location. This interpretation is further aided by the three identified PHG clusters having extensive connectivity to structures relevant to feedback processing (green, gold, and blue clusters in Figure 8). A similar set of PHG clusters was identified in an fMRI study investigating the functional specificity of local morphology of the human parahippocampal region (Huntgeburth et al. 2017). Here, the middle and posterior sections of the PHG (defined relative to the segments of the collateral sulcal complex) showed heightened activation when subjects were engaged in recalling a cognitive map to find the route to a goal location. These anatomical definitions by landmarks correspond to the pPHG, and the posterior section of the aPHG ROIs (aPHG2) from this study, which were the only three clusters predicted by RPT activity. We speculate that the clusters of activation identified in the present study may reflect these diverse PHG regions, the Parahippocampal Place Area (pPHG clusters) and the entorhinal cortex (aPHG cluster), respectively, and that enhanced synchronization of theta activity may play a role in regulating information flow within the PHG circuitry in a way that optimizes event encoding.

## General Discussion

A large body of evidence from rodent studies indicates that the PHG updates and stores contextual information related to spatial navigation according to the phase of local electrophysiological oscillations in the theta frequency range. In the present study, we provide converging evidence from EEG, MEG, and simultaneous EEG-fMRI data supporting the proposal that RPT elicited by feedback received in the virtual T-maze reflects phase resetting of the ongoing theta rhythm in the right PHG, and that the RPT phase shift between feedback presented in left and right alleys results from differences in the timing of the theta phase reset. Experiment 1 replicated the RPT effect and observed an analogous MEG component in the same group of individuals performing the virtual T-maze task. Consistent with previous work, feedback presented in the right alleys elicited an earlier and larger RPT peak, a larger ITC peak in the RPT range of 7-10 Hz, and a higher phase alignment in the MEG than left alleys. Importantly, since the MEG data yielded the same results as the EEG data and previous EEG studies, our EEG findings are likely not a result of volume conduction since MEG is less susceptible to this distortion (van den Broek et al., 1998). Moreover, with the MEG’s higher spectral resolution, we can now more confidently state that the RPT effect is in the suspected 7-10 Hz range. Crucially, in Experiment 2 this range exhibited the only positive single-trial coupling to the right PHG, strengthening the proposal that the RPT effect related to partial phase resetting is linked to the PHG.

These findings are consistent with decades of animal and computational research showing that theta phase resetting occurs across the medial temporal lobe in response to motivationally relevant spatial information (Givens, 1996; Hasselmo et al., 2009), which supports the encoding of goal-related spatial information (Burgess, 2008; Burgess et al., 2007; McCartney et al., 2004; Williams & Givens, 2003). Our findings also match those of studies examining human intracranial recordings, which report the existence of PHG cells producing navigational theta oscillations sensitive to landmarks (Ekstrom et al., 2005). In addition, our results provide further evidence for phase resetting occurring across temporal and frontal cortical areas during memory encoding and retrieval (Rizzuto et al., 2006) and, for theta power being modulated by the relevance of spatial information to goal attainment (Stangl et al., 2021). Phase resetting has been proposed to reflect a crucial neural mechanism for synchronizing oscillations by aligning their phase to relevant reference points (Canavier, 2015). For example, according to the oscillatory interference model (Burgess, 2008; Burgess et al., 2007) neural representations of spatial locations are reflected in the relative phases of theta local field potentials in the hippocampus and PHG. In this model phase resetting serves as a mechanism for establishing place cells, changing their activity when new information is discovered (i.e., rewards), and correcting error accumulation between the different oscillators. Accordingly, human phase resets in response to spatial information, as indexed by RPT, could serve to create spatial representations around motivationally relevant locations that contain rewards and coordinate this information across larger networks. This interpretation is consistent with a common account describing phase resetting and cross-frequency coupling as tools for balancing local population activity across regions of a network (Voytek & Knight, 2015). In previous studies that employed the virtual T-maze task, as well as in our present examination, analyses of EEG, MEG and fMRI data have been pointing consistently to a difference in processing of feedback stimuli in right alleys as opposed to left alleys. This was reported to occur only when navigating in the T-maze condition (Baker et al., 2015; Baker & Holroyd, 2009, 2013). Taken together, these results represent a rightward bias where the PHG responds more sensitively and earlier to information discovered following rightward trajectories. While the present results and existing literature point towards bilateral spatial processing (Ohnishi et al., 2006; Weniger et al., 2010), they are also consistent with the long-standing observation of right hemisphere dominance (Jacobs et al., 2010; Maguire, Burgess, et al., 1998; Toussaint et al., 2008; Weniger & Irle, 2006). With the current data we cannot draw any conclusions and can only speculate as to where this bias for rightward trajectories stems from. We previously proposed that during this 2D virtual T-maze task, the two goal locations in the left and right alley are represented by the same place field, since subjects remain stationary during the task and process the T-maze as a single location (M.-H. Lin et al., 2022). As a result, the two goal locations could be represented by different phase positions along the theta cycle of a single place field, which in turn resets the phase of the PHG theta oscillation at different timings. Future research will have to build on these speculations to determine their accuracy.

## Limitations

It is important to note as a limitation of this study that while most of the replications were successful, we only replicated the difference in phase alignment between left and right alleys in the EEG data of Experiment 1 with the ITC analysis but not the RVL analysis. This may be due to the far smaller sample size (n = 11) and decreased power of this experiment compared to our previous study (n = 91; Baker & Holroyd, 2013). It is likely that this phase alignment difference in the EEG would have reached significance in a larger sample, given that the spatial left and right alley difference was successfully replicated in both the EEG and MEG studies when analyzing the ITC, RPT power, and latency.

Similarly, Experiment 2 mostly replicated previous findings successfully. However, in the EEG data there was no significant RPT power difference observed between left and right alleys in the T-maze condition. Furthermore, the fMRI voxel clusters in the aPHG exhibited only weak activation for the left vs. right alley contrast. For the latter, smaller clusters with relatively small activation compared to the pPHG were in line with our previous findings, further emphasizing the dominant involvement of the pPHG in processing feedback in the virtual T-maze task. Regarding the EEG data, the lack of power difference could be attributed to the heavy artifact correction procedures due to the conditions when recording EEG inside the MRI scanner. Yet, given that the spatial distribution of RPT, its timing, as well as latency difference between left and right alleys specific to the T-maze condition were consistent with results observed in previous studies, we consider this a successful replication. Regarding the single-trial analysis and multimodal PHG parcellation, it is notable that among the PHG ROIs significantly predicted by RPT power two were in the pPHG and one was in the aPHG (i.e., aPHG2), exhibiting the same trends as the pPHG. Given this result and the overlap in connectivity between aPHG2 and posterior PHG ROIs, it is possible that aPHG2 and pPHG1 are part of the same region that were identified as unique due to individual differences between subjects and spatial smoothing.

Finally, while the current results support our hypothesis of RPT being related to theta phase resetting in the PHG, future research will have to explore the relationship between RPT and memory encoding as well as how information is processed in the hypothesized regions connected to the PHG in a larger spatial navigation network. Previously, we found that the timing of the ERP peak related to RPT and BOLD activation in the PHG during the virtual T-maze task were correlated to how well subjects recalled the shape of the maze they had navigated during the recordings (Baker et al., 2015; Baker & Holroyd, 2013). Together, these findings suggest that RPT is specific to moving in spatial environment and that RPT reflects interindividual variability in how subjects process environmental cues with respect to movement through an egocentric frame of reference. However, empirical tests of the relationship between RPT and memory encoding are still needed.

## Conclusion

The present study adds support for the proposal that feedback-related RPT power is a scalp level signature for the human PHG theta phase reset in service of goal-directed spatial navigation. Specifically, these results strengthen the idea of the PHG as the source of the human RPT effect, which has substantial implications for the clinical research and diagnostics of memory-related disorders. For example, with declines in spatial navigation abilities gaining attention as a potential screening tool for the detection of early-onset Alzheimer’s disease (Allison et al., 2019; Coughlan et al., 2018), the associated biomarkers have also garnered interest in the attempt to achieve higher diagnostic precision and sensitivity (Brueggen et al., 2017; de Toledo-Morrell et al., 2000). The present theta PHG signature has the potential to serve as a scalp-level signature of the integrity of functions like memory encoding linked to phase resetting. Future work could make use of this in guiding the diagnostic process or for non-invasive neurostimulation interventions in memory-related disorders, such as Alzheimer’s disease.

## Methods

### Experiment 1: Asynchronous EEG-MEG

#### Participants

Twelve participants (MAge = 25 ± 2.76 years; 5 female) participated in Experiment 1. One subject was excluded due to excessive artifacts during the MEG recordings (MAge = 25 ± 2.9 years; 5 female). All participants had normal or corrected-to-normal vision and none reported a history of head injuries. Participants were graduate students recruited from the University of Victoria (BC, Canada) and were compensated C$50 for completing the study, plus a monetary bonus associated with the experimental task. Participants also received a monetary bonus equivalent to the amount of rewards they found in the T-maze task. All participants gave informed consent. The study was approved by the University of Victoria research ethics committee and was conducted in accordance with the ethical standards described in the 1964 Declaration of Helsinki.

#### Virtual T-maze Task

For the EEG and MEG experiments, subjects performed the virtual T-maze task (Figure 1) which had also been used in our previous EEG studies (Baker et al., 2020; Baker & Holroyd, 2009, 2013). In this task, subjects were instructed to maximize their rewards (five cents) by going either the left or right in the virtual T-maze, which consisted of still images depicting cardinal locations in the maze. On each trial subjects started out in the stem of the maze looking down towards a junction point where two alleys branch off orthogonally from the stem. After 1000 ms, subjects were presented with a green double arrow prompting them to choose the left (button 1, left index finger) or right alley (button 2, right index finger). As soon as a valid response was given, they moved to the junction point looking down the right or the left alley for 1000 ms and received feedback to their choice for another 1000 ms (a still image of an apple for reward or a still image of an orange for no reward). Unbeknownst to the participants, the type of feedback was selected at random (50% probability). The complete task consisted of four blocks with 100 trials each. At the end of the task, subjects received the total amount of rewards as a monetary bonus.

In the behavioral domain the sum of choices (left vs. right), reaction times, and response strategies were analyzed to a) ensure that there were no differences in how subjects responded during left and right trials and b) to test if subjects responded to feedback in a similar manner as in previous studies. The full results of the behavioral analysis are detailed in Figure 1-figure supplement 1.

#### EEG and MEG data acquisition

The EEG data were recorded using a montage of 63 electrode sites in accordance with the extended international 10-20 system (Jasper, 1958). Signals were acquired using Ag/AgCl ring electrodes mounted in a nylon electrode cap with an abrasive, conductive gel (Falk Minow Services, Herrsching). They were amplified by low-noise electrode differential amplifiers with a frequency response of DC 0.017-67.5 Hz (90 dB octave roll off) and digitized at a rate of 250 samples per second. Digitized signals were recorded to disk using Brain Vision Recorder software (Brain Products GmbH, Gilching, Germany). Inter-electrode impedances were maintained below 10 kΩ. Two electrodes were also placed on the left and right mastoids. The EEG was recorded using the average reference. The electrooculogram (EOG) was recorded for the purpose of ocular correction; horizontal EOG was recorded from the external canthi of both eyes, and vertical EOG was recorded from the suborbit of the right eye and electrode channel Fp2.

MEG data were acquired using a 151-channel Canadian Thin Films (CTF MEG, Coquitlam, BC, Canada) whole-head magnetometer with axial first order gradiometers (VSM MedTech). Four EEG (Cz, Fz, left mastoid, right mastoid) and two EOG channels were added, one horizontal EOG channel on the external canthi and one vertical EOG channel in the suborbit of the right eye. Participants sat in an upright position while performing their task, which was projected onto a screen in front of them. Data were recorded continuously for approximately 15 minutes and digitized at 600 Hz. To ensure a standardized co-registration of sensors and the head, fiducial head positioning coils were arranged around the nasion and the bilateral preauricular points before the experiment.

#### EEG and MEG data preprocessing

Both EEG and MEG analyses were performed in Python 3.9 and MNE-python (Version 1.0.0) for M/EEG analysis (Gramfort et al., 2014). Data were bandpass filtered (low-pass: 60 Hz, high-pass: 0.1 Hz) to remove power line noise, other high frequency noise and slow, movement induced signals. To remove ocular and movement artifacts, data were entered into an Independent Component Analysis (ICA) and decomposed into independent components (IC) using the Infomax algorithm. The ICA was run on a copy of the data, which was high-pass filtered at 1 Hz to prevent distortion through high amplitude, low frequency signals (Winkler et al., 2015). Next, filtered EOG signals were segmented around likely saccades and blinks. For the MEG, segments around likely heartbeat artifacts marked by their sharp, large peaks surrounded by ripples mirroring the QRS-complex were extracted as well. The time courses of each component were correlated to these ocular and cardiac artifact segments in order to identify components representing artifact signals. Furthermore, potential artifact components were inspected based on their topography and spectral composition. Components marked as artifacts were excluded from back-projection to continuous data. The cleaned EEG data were re-referenced to the bilateral mastoids. Lastly, all data were segmented around feedback presentation (±2500 ms). Segments exceeding a maximum peak-to-peak amplitude of 100 μV were marked and excluded from the time-frequency analysis. For the MEG the peak-to-peak amplitude threshold for magnetometers was set to 2000 fT.

#### EEG and MEG time-frequency analysis

For the time-frequency analysis of all EEG and MEG data, cleaned segments time-locked to feedback presentation (±2500 ms) and separated by maze context and alley, were exported to MATLAB (release 2020b, Mathworks, Massachusetts, USA) and analyzed using custom MATLAB scripts from our previous study (Baker & Holroyd, 2013). The frequencies ranging from 1-50 Hz were analyzed using a complex seven-cycle Morlet wavelet for convolution. Total spectral power was obtained by averaging the EEG spectrum across all trials and time for each subject. Time-frequency analysis on the single-trial EEG data thus yielded total theta power, including theta power that was both phase consistent (evoked) and phase inconsistent (induced) across trials with respect to the eliciting event. Evoked theta power was determined directly from the averaged ERPs, and induced theta power was then identified by subtracting the evoked theta power from the total theta power (for more details, see Baker & Holroyd (2013); Hajihosseini & Holroyd (2013)). The relative change in the power for each condition was determined by averaging the baseline activity (-200 to -100 ms prestimulus) across time for each frequency and then subtracting the average from each data point following stimulus presentation for the corresponding frequency. This value was then divided by the baseline activity to normalize the change of power to the baseline activity. To note, spectral power was reported here unitless because it is calculated as a proportional increase/decrease relative to baseline. For each subject and feedback location, we identified the maximal increase in power for each subject’s peak theta frequency within the range from 4 to 12 Hz. The peak power and latency of each frequency band were obtained by detecting the maximum power within a 600 ms window following the onset of the feedback stimulus. This was done separately for right and left alleys.

#### EEG and MEG intertrial coherence and phase analysis

The analysis of phase resetting relied on two main parameters: 1) inter-trial coherence (ITC) as a measure of correlation between phase angles across individual trials, as well as 2) the mean phase degree angle and spread of phase. To establish phase resetting, phase coherence for the theta band after feedback across trials should increase across trials compared to baseline. Then, to determine if this coherence is indicative of a reset, the spread of phase angles should decrease due to phase alignment. This analysis was performed on the same five seconds of time-domain data centered around feedback presentation (±2500 ms) as used for the time-frequency analysis. To match our previous analysis (Baker & Holroyd, 2013), the segments were trimmed to a time window from 500 ms pre-feedback to 1000 ms post-feedback. For the ITC analysis, we identified the frequency with maximum power in the EEG and MEG data for this time window. Then the algorithm implemented in the pop_newtimef function in EEGLAB (v2022, Delorme & Makeig, 2004; Makeig et al., 2002) assigns to that frequency a value between 0 and 1, where 1 indicates perfect EEG phase consistency across trials and 0 indicates random EEG phase distribution across trials. Lastly, the peak ITC value and latency were extracted within the general time window of the RPT effect. For the phase analysis, phases (in degrees) at frequencies in multiples of 2 Hz (i.e., 5-6 Hz, 7-8 Hz, 9-10 Hz, and 11-12 Hz) were extracted in a 10 ms window centered around the mean peak timing of the RPT effect averaged across alleys (EEG: 186 ms, MEG: 196 ms). In addition, the baseline window (i.e., -186 ms, -196 ms) was also analyzed as a control time period. Phase data was extracted for each trial, alley, and subject at PO8 for the EEG, as well as for each subject’s peak channel from the right parietooccipital MEG sensor cluster identified in the time-frequency analysis. Differences in ITC latency and strength between left and right alleys were tested using one-sided Wilcoxon signed-rank tests.

In order to determine the circular mean of phase angles across trials, alleys, and subjects, the CGM and the RVL were examined, using the Circstat MATLAB toolbox (Berens, 2009). These two measures served to assess differences in mean angle as well as spread of phase angles. The CGM represents an arithmetic mean for a circular scale and, thus, provides information on which phase angle, on average, occurred across trials and subjects for a particular frequency. How strongly individual phase angles spread around this mean orientation is captured by the RVL, where a value close to zero indicates no alignment and a value close to one indicates ideal alignment around the mean phase direction. Hence, the RVL was used as the main indicator of phase alignment. In order to test differences in the CGM between conditions, the Harrison-Kanji test (Harrison & Kanji, 1988) as well as the Watson-Williams test (G. S. Watson & Williams, 1956) were employed. These tests are circular analogues of a two-factor and a one-factor ANOVA respectively. The one-factor variant was used to determine if there was any difference between the pre- and post-feedback window. The two-factor test assessed the effects of the factors Feedback Alley Type (left vs. right alley) and Frequency (8 Hz vs. 10 Hz) as well as their interaction. For the RVL, an analogous conventional two-way repeated measures ANOVA was used. This analysis was run using the same custom MATLAB and EEGLAB scripts as in Baker and Holroyd (2013). Post-hoc tests between factor levels were performed using one-sided Wilcoxon signed-rank tests.

### Data availability

Analysis scripts, documentations, and examples for both preprocessing and higher level statistics can be found in the folders for Experiment 1 in the github repository for this study: https://github.com/BakerlabRutgers/phg_phase_resetting. All further code and data are available upon reasonable request.

## Experiment 2: Simultaneous EEG-fMRI Participants

For Experiment 2, 28 right-handed undergraduate students (MAge = 23.5 years, SDAge = 4.24 years; 16 female) were recruited from Rutgers University-Newark and the New Jersey Institute of Technology (NJ, USA). Three of these subjects were excluded due to a lack of valid trials. The resulting sample had a mean age of 23.6 years (SDAge = 4.31 years; 15 female). All subjects were screened with self-report questionnaires regarding demographic information, handedness (Oldfield, 1971), spatial navigation abilities (Hegarty et al., 2002), neurological and psychiatric history, their vision, and MRI permissibility (i.e., pregnancy, metal or medical implants, etc.). Only subjects who were right-handed, between 18 and 55 years of age, had normal-to-corrected vision, and had no history of neurological or psychiatric disorders or medication were allowed to participate. For their participation, subjects received course credit as well as monetary compensation equivalent to the amount of rewards they earned during the task. The study was approved by the Rutgers University Ethics Committee Board and was conducted in accordance with the ethical standards described in the 1964 Declaration of Helsinki.

### Maze/No-maze Task

Participants in the simultaneous EEG-fMRI experiment engaged in the same version of the Maze/No-maze task (Figure 5) used in our previous fMRI study (Baker et al., 2015). In this task, 192 trials were divided equally across six blocks with each trial following the same progression of events as in the virtual T-maze task (Figure 1B). However, blocks alternated between the T-maze and No-maze condition in a counterbalanced manner across subjects. In addition, rest trials displaying a fixation cross for 3000 ms were randomly dispersed across blocks (16 per block) to optimize fMRI design efficiency. In both conditions subjects performed the same T-maze task as described before, except that in the No-maze condition there was no maze background. Instead, response prompts and feedback images were shown on scrambled versions of the maze backgrounds. As a result, the No-maze condition had no spatial context. Behavioral data, demonstrating the equivalency of left and right alleys (sum of choices, reaction times, and response strategy) are described in Figure 5-figure supplement 1.

### EEG-fMRI data acquisition

EEG data for all 256 channels were collected at a sampling rate of 1 kHz using an MR-compatible HydroCel Geodesic sensor net (Electrical Geodesics Inc., Eugene, USA). In addition to scalp electrodes, a four-lead electrokardiogram (EKG; two active, two dummy) was recorded for later artifact correction. One active EKG lead was placed on the lower end of the sternum and one below the left chest on the rib cage. The amplifier was shielded in a Field Isolation Containment System (FICS), allowing it to be placed on the ground next to the bore. All wires were carefully straightened beside the subject to prevent hazardous wire loops. The FICS was connected via a fiber optic cable to an intermediary USB clock sync interface in the control room. At the USB interface, digital inputs from the experimental computer and the MRI clock, event triggers of the Maze/No-maze task, and MRI triggers for volume acquisition were relayed to the EEG acquisition computer. This enabled high precision synchronization of EEG and MRI acquisition necessary for MR-related artifact correction. All electrodes recorded from inside the scanner were online referenced to E257, which is the geodesic electrode placement system’s equivalent to Cz in the international 10-20 system (Jasper, 1958). Conversions of electrode positions in the geodesic sensor net to the nearest equivalent in the 10-10 system were based on Luu & Ferree (2000). Data were recorded using the Net Station software (Version 5.4.2) applying an online bandpass filter excluding data above 100 Hz and below 0.001 Hz. All electrode impedances were kept below 50 kΩ.

MRI data were collected in a 3T scanner (Trio Tim, Siemens) with a 12-channel head coil at the Rutgers Brain Imaging Center. Anatomical images were acquired with a T1-MPRAGE sequence, consisting of 176 sagittal slices (1 mm isotropic voxel; TR = 2500 ms, TE = 2.52 ms, flip angle = 9°). A dual-echo gradient-echo sequence was used to assess a B0 inhomogeneity gradient field map (TE1 = 5.19 ms, TE2 = 7.65 ms, TR = 400 ms). Functional whole brain images were collected in an axial orientation by using a T2-weighted gradient echo planar imaging sequence (TR = 2000 ms, TE = 25 ms, flip angle = 90°) with an isotropic voxel size of 3.3 mm (64 x 64 matrix; 208 x 208 mm2 field of view). 35 slices per volume were acquired in an ascending, interleaved order. Functional image acquisition sessions lasted 15 minutes, resulting in 450 volumes for each participant. Moreover, a resting state and diffusion weighted imaging acquisition was added after the functional task runs, lasting approximately 12 minutes each.

### EEG data preprocessing and analysis

EEG data concurrently collected with fMRI are affected by two major MRI-related artifacts: Gradient artifacts (GA) and ballistocardiac artifacts (BCA). Both GA and initial BCA correction based on principle component analysis were carried out using Brain Vision Analyzer 2.1 (Brain Products GmbH, Gilching, Germany). Subsequent preprocessing including ICA, filtering, re-referencing, and segmentation were performed using custom scripts in Python 3.9 and MNE-python (Version 1.0.0) for M/EEG analysis (Gramfort et al., 2014).

The first step in cleaning the EEG data was to subtract GAs caused by the periodically changing magnetic fields during the acquisition of functional images. For this purpose, an average artifact correction (Allen et al., 2000; Moosmann et al., 2009) was performed. An artifact template was built from the EEG activity in 25 GA windows around a given artifact (±12 GAs). This template was subtracted by moving it across the continuous signal contaminated by GAs. To account for BCAs, a peak detection was run to find R peaks in the EKG channel. Results of this detection were inspected and manually corrected if needed. Using the first three components from a principle component analysis run on the heartbeat segments centered around the R peaks in the EKG, a new BCA template was formed and subtracted from the EEG signal (Niazy et al., 2005).

Next, the EEG signals were filtered using a Butterworth filter with a bandpass of 0.1-60 Hz. To remove residual BCAs as well as ocular and movement artifacts, all channels were entered into an ICA and decomposed into independent components using the Infomax algorithm. As in Experiment 1, a copy of the data was high-pass filtered at 1 Hz before running the ICA on this copy. When unmixing the EEG signal, 60 ICA components were extracted for each subject. BCAs are primarily caused by movement of electrodes and cardiac-related activity (Allen et al., 1998). Due to the pulsatile blood flow at temporal arteries while lying in the scanner, they occur as a prototypical progression of topographies marked by reversed polarities over the left and right hemisphere and higher amplitudes at temporal electrodes (Yan et al., 2010). Another reliable indicator of BCAs is their time course correlation to the ejection phase of the cardiac cycle, as indexed by the EKG signal’s R peak, and more broadly the QRS complex, which marks a rapid depolarization of the right and left heart ventricles (Debener et al., 2008; Shams et al., 2015). BCA components were thus identified based on their topography and Pearson correlation to the EKG segments. Components reliably categorized as BCAs at the end of this validation procedure as well as components reflecting eye or muscle movement were rejected before back-projecting the ICA to the continuous EEG signal.

After data were both GA- and BCA-corrected, they were re-referenced to the average of all scalp electrodes, excluding sensor net positions located on the cheeks or the neck. The continuous data were segmented into five second windows around the presentation of positive and negative feedback (±2500 ms). This was done separately for both T-maze vs. No-maze trials as well as feedback presented in left and right alleys. After all preprocessing was completed, three out of the 28 original subjects were excluded due to a lack of valid segments. For the remaining 25 subjects, the same time-frequency analysis as described for Experiment 1 was performed on the geodesic equivalents of PO7 (E107) and PO8 (E160). For each subject, 4-12 Hz power was averaged across all conditions. Then, the peak theta frequency for each subject and condition was determined.

Peak RPT power and latency values (50 ms to 350 ms) were entered as dependent variables into two separate two-way repeated measures ANOVAs. Both tested the factors Feedback Alley Type (left vs. right alley), Task Type (Maze vs. No-maze) as well as their interaction. To ensure homogeneity of variance a Shapiro-Wilk test was run and approximate normal distribution of the dependent variable was tested using a Levene test. Post-hoc multiple pairwise comparisons of main effects and simple effects of the interaction were run using paired t-tests. Corresponding p-values were corrected using the Bonferroni-Holm method.

### fMRI data preprocessing and analysis

MRI data were formatted according to the international Brain Imaging Data Structure (BIDS) (Gorgolewski et al., 2016). Apart from this, the preprocessing of the functional data was performed identically to the previous study (Baker et al., 2015) using custom scripts in MATLAB (release 2020b, Mathworks, Massachusetts, USA) and SPM12. Functional data were first slice-timing and then motion corrected with respect to the first image. Afterwards, functional images were co-registered with the structural scan and then normalized to a standard Montreal Neurological Institute (MNI) template with 12-parameter affine registration. All normalized images were smoothed with an 8-mm^3^ FWHM Gaussian kernel.

First-level analyses were performed using a general linear model (GLM) including a constant term, six motion regressors obtained from motion correction, and eight event regressors to model activation time-locked to feedback presentation as well as orientations for the T-maze and No-maze conditions. Events were divided by feedback valence and alley. These four events were further divided by separating by T-maze and No-maze conditions, resulting in four regressors for each alley and feedback. These were built identically to previous experiments using the fMRI version of the Maze/No-maze task (Baker et al., 2015). All models were run with a canonical hemodynamic response function and its temporal derivative. Before the preprocessed time series data were statistically modeled, they were high-pass filtered with a mean cutoff period of 128 s. To identify voxel activation related to spatial effects, all feedback presented in the T-maze condition was contrasted to feedback presented in the No-maze condition. To identify activation associated with navigating towards the left or right, directions were contrasted separately for T-maze and No-maze conditions. Finally, linear contrasts of coefficients for each participant were used to run a second level analysis by applying paired t-tests and FDR correction for multiple comparisons.

As described in the results section of Experiment 2, we constructed a PHG mask consisting of twelve bilateral ROIs based on connectivity patterns found in DWI (Baker, Reid, et al., 2017) data from the HCP (Van Essen et al., 2012). This approach was used to extract regional variations in the connectivity profile of the selected seed region with the whole brain. To note, the boundaries of the aPHG and pPHG region were based on the definition described by Huntgeburth and Petrides (2012). The tractography-based parcellation results yielded four subregions within the right aPHG, and two subregions within the right pPHG (Figure 8B) (see section below).

Standardized beta series values were then extracted from the twelve bilateral PHG clusters (six per hemisphere; Figure 8B) using nilearn NiftiMasker (Abraham et al., 2014). These time series were created by splitting regressors using the BIDS application NiBetaSeries for python (Kent & Herholz, 2019). Instead of a conventional GLM with one beta for each event regressor, beta coefficients were estimated for every single event in the design matrix. To counteract the overlap of BOLD responses on single-trials, we employed the least squares separate (LSS) approach (Mumford et al., 2012; Rissman et al., 2004). LSS improves the estimation of single-trial event-related BOLD signals with short intervals between events (i.e., 3-4 seconds) by calculating one GLM for each trial with one predictor for the trial of interest and one for the combination of all the other trials.

### PHG segmentation analysis: Diffusion-weighted and resting-state functional MRI

*Participants and data acquisition* We investigated the anatomical and functional organization of the human PHG using diffusion and functional MRI data from the HCP. Data were obtained from the 500-subject release of the HCP (HCP RRID:SCR_008749) database from March 2015. The multimodal data used here include structural MRI, diffusion-weighted MRI, and resting-state MRI. The scanning procedures are described in detail in Van Essen et al. (2012) and are available online (https://www.humanconnectome.org). In total, 430 subjects’ data were preprocessed and passed quality control.

*Seed regions* The seeds of the anterior and posterior PHG regions were defined manually by reference to the morphology of the sulci of the collateral sulcal complex as described by Huntgeburth and Petrides (2012) on an average normalized (ICBM152) anatomical scan using the DISPLAY software package (MacDonald, 1996). In brief, the posterior boundary of the PHG, and thus the parahippocampal cortex, is based on the junction between the collateral sulcus proper and the PHG extension of the collateral sulcus. The point where the PHG extension and the collateral sulcus proper merge provides the point of origin of the occipital extension of the collateral sulcus which continues into the lingual gyrus, thereby forming the posterior limit of the PHG. In regards to the anterior boundary of the PHG, the rhinal sulcus forms the lateral limit of the entorhinal cortex that is located on the anterior portion of the PHG, with the caudal end of the rhinal sulcus marking the posterior border of the entorhinal cortex. The PHG extends posterior to the entorhinal cortex and, therefore, the rostral origin of the anterior segment of the collateral sulcus proper provides an anatomical marker of the anterior limit of the PHG. Furthermore, the collateral sulcus proper forms the lateral boundary of the PHG, delimiting it from the adjacent fusiform gyrus. When the collateral sulcus proper can be divided into two sulcal segments, the transition between the anterior and posterior segments occurs approximately at a y-coordinate of -33 in each hemisphere. The two segments of the collateral sulcus proper thus form anatomical landmarks that may be used to differentiate between the aPHG versus the pPHG: the aPHG lying medially along the anterior segment of the collateral sulcus proper, and the pPHG along the posterior segment of the collateral sulcus proper, rostral to the lingual gyrus. Once defined, the aPHG and pPHG masks were linearly registered to native diffusion space.

*Diffusion MRI* Diffusion data were collected with 1.25 mm isotropic spatial resolution and three diffusion weightings using HCP dMRI protocol (Sotiropoulos et al., 2013) and downloaded using the HCP Diffusion pipeline (Glasser et al., 2013). The probability distributions of fiber orientation were estimated by using FSL’s (RRID:SCR_002823) multi-shell spherical deconvolution toolbox (bedpostx), where each voxel contains at most three fiber directions and the diffusion coefficients were modelled using a Gamma distribution (Jbabdi et al., 2012). A T1- weighted image downsampled to the resolution of the diffusion data was employed for the nonlinear registration of the anterior and posterior seed from MNI standard space to native structural volume space using FSL’s package FNIRT. The parcellation of aPHG and pPHG was carried out on 40 randomly selected subjects (20 female, 20 male) in order to limit computation time and data storage. To test the robustness of parcellation, we randomly divided the 40-subjects dataset in half and applied the parcellation procedure (as explained below) independently to each group. The first 20 subjects were used as the test group to reveal the underlying organizational pattern of the PHG and a second group of 20 subjects was subsequently used as the replication group to test the stability of our parcellation maps. All diffusion data (n = 430) were then used during the tractography analysis to map the connectivity profiles of each of the PHG subdivisions identified by the parcellation (please see Zhang et al. (2017) for full details).

*Connectivity-based parcellation of PHG* A data-driven connectivity-based brain parcellation procedure was used as described in Zhang et al. (2017) (see also Fan et al., (2014, 2016)). First, probabilistic tractography was applied by sampling 5000 streamlines at each voxel within the aPHG and pPHG seed mask. A target mask was constructed for each subject that included all brain voxels (white or gray matter) connecting to the seed region. The whole brain connectivity profile for each aPHG and pPHG voxel was then saved as a connectivity map and used to generate a connectivity matrix with each row representing the whole brain connectivity profile of one seed voxel. Next, a correlation matrix was calculated as a measure of similarity between the connectivity profiles of each voxel pair with the aPHG and pPHG (Johansen-Berg et al., 2004). Spectral clustering was applied to the similarity matrix to identify clusters with distinct connectivity profiles (H. Liu et al., 2013; Shi & Malik, 2000). We applied this procedure separately for each subject and each hemisphere to generate a series of parcellation maps for all individuals at different resolutions (i.e. numbers of regions/parcels) and chose cluster numbers ranging from 2 to 6 in each hemisphere, subsequently using the most stable and consistent parcellation map. The optimum parcellation solution (i.e. number of parcels) was then determined by evaluating the reproducibility of parcellation maps through a split-half procedure. More specifically, we randomly split the entire group 100 times into two non-overlapping subgroups and generated the group parcellation maps for each subgroup separately. The consistency between each pair of parcellation maps was evaluated by using the normalized mutual information (P. Zhang, 2015). The average indices among 100 samples were calculated to represent the stability of each parcellation. The suitable cluster number was then determined by searching for the local peaks in the stability curve.

*Connectivity profile of each PHG subdivision* Based on the obtained parcellation map of the aPHG and pPHG, we mapped the anatomical connectivity profiles of each subdivision by performing probabilistic tractography with 10,000 streamlines from each PHG subdivision. The resulting connectivity maps were first normalized by the size of the seed region and total number of streamlines (i.e. 10,000) in order to generate the relative tracing strength from the seed to the rest of the brain. A threshold of 0.001 (i.e. 10 out of 10,000) was then used to remove noise effects of fiber tracking. The resulting individual tractograms were combined to generate a population map of the major fiber projections for each aPHG and pPHG subdivision. Another probabilistic threshold of 50% was applied to the population fiber-tract maps (i.e. at least half of subjects showing each retained fiber tract). This resulted in a group averaged tractogram for each subdivision of aPHG and pPHG. Finally, a maximum probability map (MPM) of fiber tracts, which represented distinct components of fiber projections for each subdivision, was also generated based on the population fiber-tract maps. Specifically, a connectome mask was first generated for each subdivision by binarizing its group tractography map with connectivity probability at 0.001. Each voxel within the combined connectome mask was then classified according to the PHG subdivisions with which it had the highest connectivity. This calculation of a MPM on probabilistic tractography has been widely used in subdividing brain structures, including the thalamus (Behrens et al., 2003), amygdala (Saygin et al., 2011), striatum (M. X. Cohen et al., 2009), and substantia Nigra (Y. Zhang et al., 2017). Here, we use this method to generate the organizational topography of fiber projections among anterior and posterior subdivisions.

*Functional connectivity patterns of the aPHG and pPHG subregions* The data used here was the 430 participant subset from the “500 Subjects” HCP release. Preprocessing consisted of standard resting-state functional connectivity preprocessing. Resting-state data collection details for this data set can be found elsewhere. Spatial normalization to a standard template, motion correction, and intensity normalization were already implemented as part of the HCP in a minimally processed version of the data set described elsewhere. With the volume version of the minimally preprocessed data, we used AFNI58 to additionally remove nuisance time series (motion, ventricle and white matter signals, along with their derivatives) using linear regression, to remove the linear trend for each run and spatially smooth the data. The data were smoothed using a non-Gaussian filter (nearest neighbor averaging) at 4 mm. For the main analyses, we used the aPHG and pPHG subregions derived from the connectivity-based parcellation of PHG for the functional connectivity seed regions. Additional regions included two reinforcement-related frontal targets (anterior cingulate cortex and ventromedial prefrontal cortex) and two spatial-related posterior targets (middle occipital cortex, precuneus) derived from our fMRI (Figure 7). Analyses were carried out with Matlab 2014b (Mathworks, Massachusetts, USA).

*Functional connectivity estimation.* The initial analyses estimated functional connectivity (FC) using Pearson correlations between time series (voxels within each PHG subregion) from all voxels of brain regions. All computations used Fisher’s z-transformed values, which were reconverted to r-values for reporting purposes. Resting-state fMRI time series from all other regions were used as predictors of the to-be-predicted region’s resting-state fMRI time series. The resulting betas, which were directional from the predictor regions to the predicted region, were then used as FC estimates. Note that beta estimate directionality reflects optimal linear scaling of the source time series to best match the target time series (based on resting-state fMRI data).

*RSFC Statistics*. All statistical inferences with empirical fMRI data that produced p-values were made using two-tailed one-sample t-tests relative to 0 (n = 430; degrees of freedom: 429). Pearson correlation (r) was used as a measure of pattern similarity, with p-values only calculated for group-level inferences using two-sided one-sample t-tests on the Fisher’s z-transformed r-values. We used a FDR correction for multiple comparisons across all calculated p-values reported in this study. This revealed an uncorrected p < 0.004 threshold. All p-values reported as statistically significant were below this threshold, such that all significant p-values were statistically significant (p < 0.05) after correcting for multiple comparisons across all analyses. For presentation purposes, only p-values below 0.00001 were reported as p < 0.00001 based on the convention.

### Integrated EEG-fMRI analysis

To test whether EEG and fMRI data were significantly coupled on a trial-by-trial level, a series of LME models were estimated, one for each of the twelve PHG ROIs. This was done separately for each condition (T-maze vs. No-maze) using four centered EEG regressors (delta: 1- 4 Hz, theta: 5-6 Hz, 7-8 Hz, 9-10 Hz, 11-12 Hz) and a random intercept for each subject. Each of these EEG regressors received a fixed effects estimate (regression weight) which was transformed to a standardized beta weight and tested for significance with a t-test. P-values within each model were corrected for multiple comparisons using the FDR method. Building on this PHG analysis, we conducted a control analysis across the whole brain by running one LME with the same four EEG regressors for each ROI in the Glasser atlas split across the left and right hemisphere (Glasser et al., 2016).

All analyses were carried out using the lme4 package (Bates et al., 2015) in the R Programming Environment (R Development Core Team, 2016). Since single-trial regressors reflect a direct co-variation of two continuous variables, such as EEG power and BOLD beta series values, LMEs allowed insight into trial-by-trial co-variation. LMEs also provide the advantage of modeling single-trial variation within single subjects and to group error terms more flexibly. Here, random intercepts were included to better group individual variance for each subject. This approach has previously been successfully applied in a simultaneous EEG-fMRI study linking single-trial visually evoked gamma activity to visual cortex activation (Beldzik et al., 2019).

The LME models tested whether centered EEG regressors derived from single-trial means of evoked power (50 ms to 250 ms post-stimulus) could predict variation of beta series values from the PHG ROIs. Due to increased noise on a single-trial level, each trial was baseline corrected with the mean of the baseline window (-200 to -100 ms pre-stimulus) averaged across all trials. BOLD activation was predicted using power values recorded at the right hemispheric EEG sensor with the highest average RPT power (E160). For each of these sensors, four EEG-based regressors were built, one for each analyzed frequency band.

### Data availability

Same as for Experiment 1, analysis scripts as well as the twelve bilateral PHG ROIs as a binary nifti file can be found in the github repository for this study: https://github.com/BakerlabRutgers/phg_phase_resetting. All further code and data are available upon reasonable request.

## Acknowledgements

We thank G. Karpov, P. Shafeek, K. Biernacki, M.-H. Lin, J. Stringfellow, and E. Winfield for assistance with the data collection for Experiment 2.

## Author contributions

TEB and MRG conceptualized the study. Experiment 1 (EEG-MEG): TEB, KK, and CH designed the study, TEB collected the data, and MRG analyzed the data. Experiment 2 (EEG-fMRI): TEB, MRG, and RDM designed the study protocol, and MRG collected and analyzed the data. TEB, YZ, AR, SCH, MP and AD created the diffusion-based parahippocampal ROIs, and MWC and TEB analyzed the resting-state functional-connectivity MRI data. MRG and TEB wrote the paper with suggestions by all authors.

## Supplementary Material

**Figure 1-figure supplement 1.**
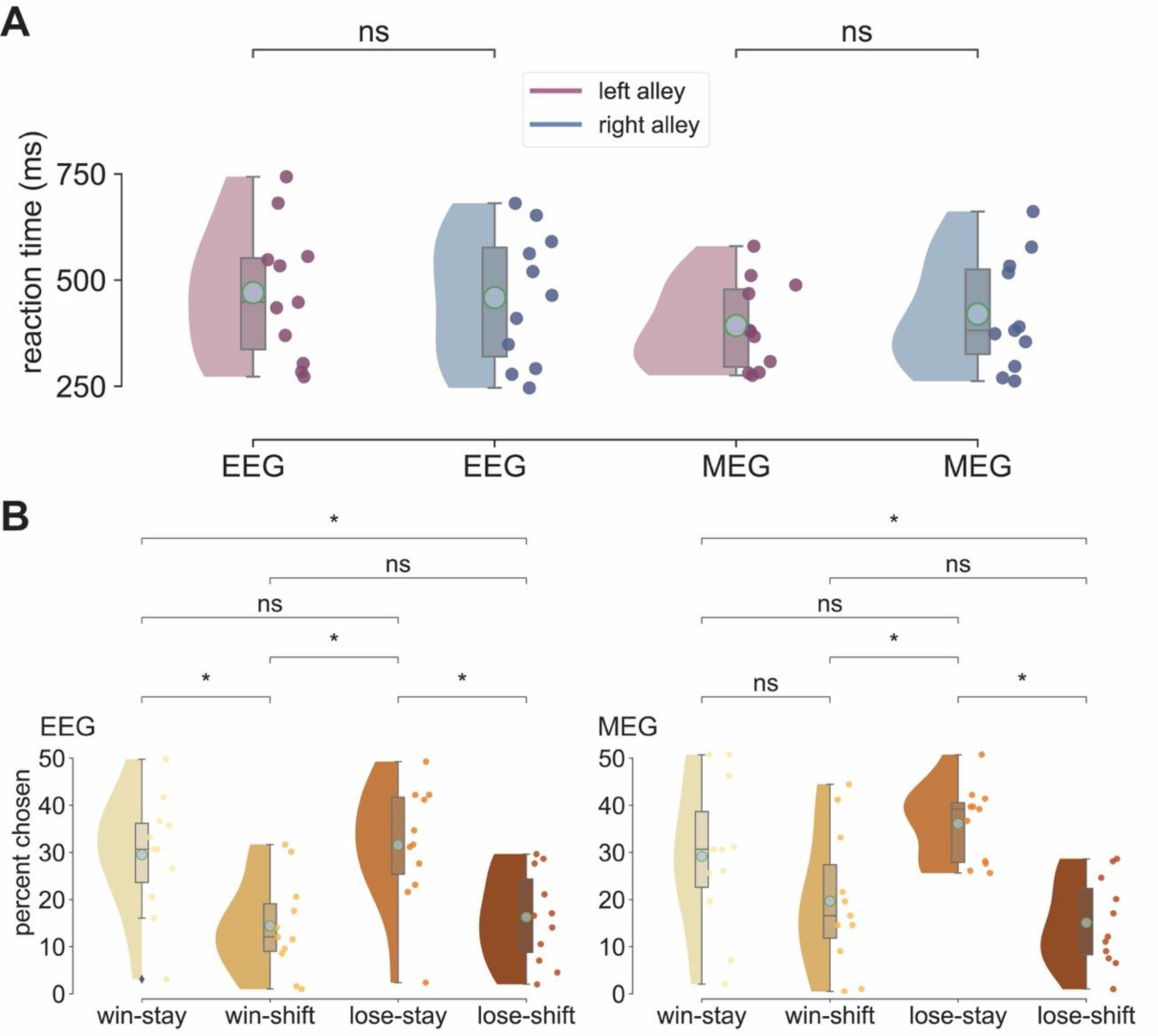
Experiment 1: Behavioral data analysis. **A**: Raincloud plots show RT density distributions with boxplots and individual subject averages in milliseconds (ms) plotted separately for each Feedback Alley Type (left in pink, right in blue) and modality. Cyan dots mark the mean value of each distribution. As has been reported before, right and left alleys in both EEG and MEG were behaviorally equivalent. In the EEG session, there was no significant difference between alleys in regard to how frequent either alley was chosen (*W* = 31, *p* = 0.859). In regard to reaction times, right alleys (M = 459 ms, SD = 147 ms) were chosen at approximately the same speed as left alleys (M = 470 ms, SD = 151 ms), *W* = 17, *p* = 0.154. During the MEG session, subjects also did not choose either alleys more frequently than the other (*W* = 19, *p* = 0.213) and responded with approximately the same reaction times for right (M = 420 ms, SD = 127 ms) as for left alleys (M = 392 ms, SD = 100 ms), *W* = 13, *p* = 0.075. **B**: Percentage of response strategies relative to total choices. Same as in A, cyan dots are the mean of each distribution and dots colored like the density plots are each individual subject’s mean value. In both the EEG and MEG session, the two most prevalent patterns were win-stay and lose-stay, each accounting for roughly 30% of total responses. In the EEG session, there was a significant main effect for the factor response strategy, *Q* = 16.1, *p* = 0.001. Consistent with previous experiments, post-hoc tests showed that win-stay when compared to win-shift was significantly more frequent (*W* = 6, *p* = 0.016). However, unlike in previous experiments, lose-shift was a significantly more common response strategy than lose-stay (*W* = 5, *p* = 0.022), indicating a tendency for sticking to the same alley regardless of feedback. A similar pattern of results was observed for the MEG session, with a significant main effect for response strategy (*Q* = 10.96, *p* = 0.012) and win-shift and lose-stay as the more common response strategies. Here, win-stay and win-shift were not significantly different (*W* = 19, *p* = 0.213), but lose-stay was significantly more frequent than lose-shift (*W* = 3, *p* = 0.008). *ns: not significant, *p < 0.05*

**Figure 2-figure supplement 1:**
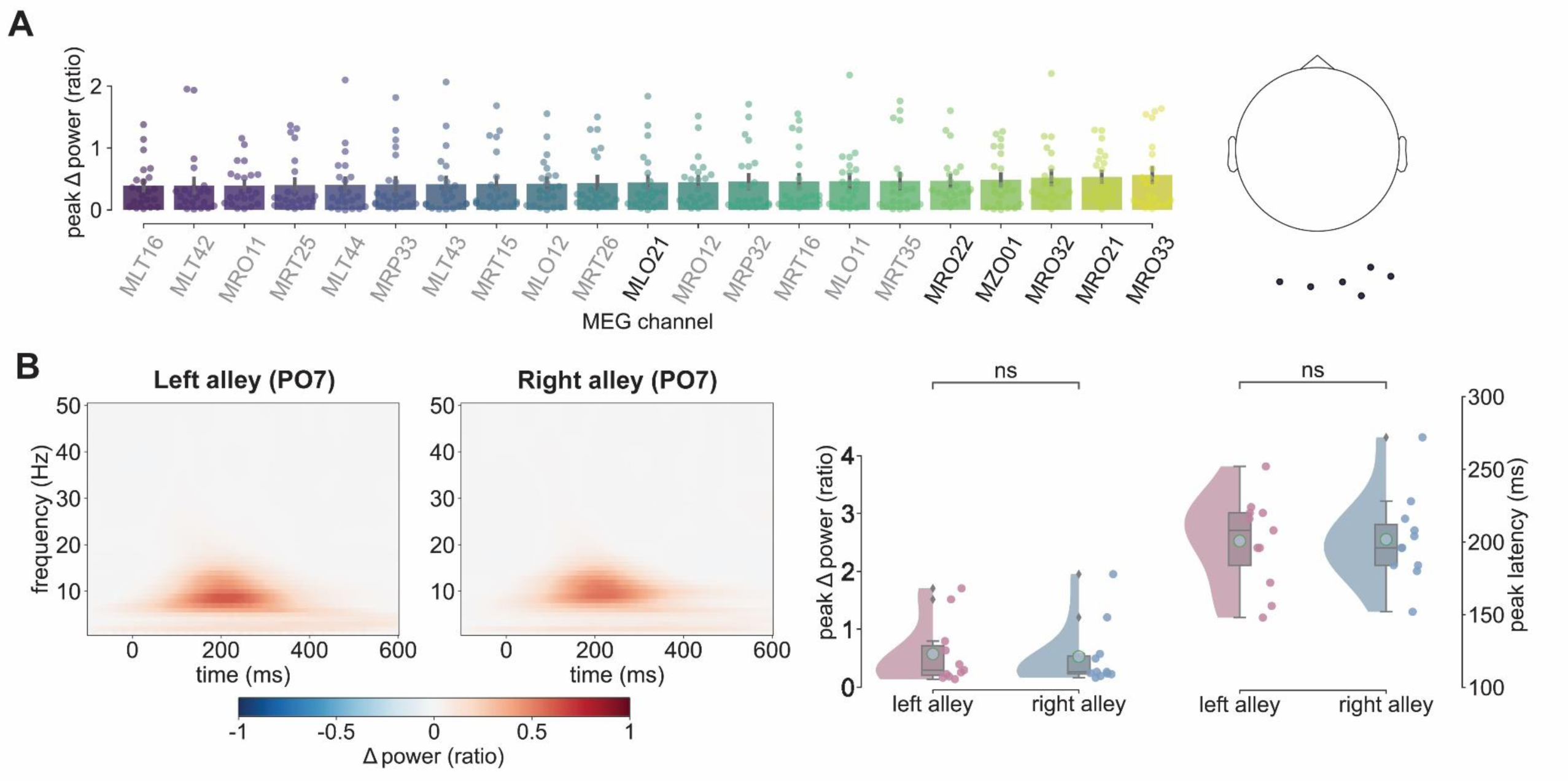
MEG peak RPT channel statistics and EEG left hemisphere RPT results. **A**: Bar plots highlight the 20 MEG channels with the highest peak evoked RPT power post-feedback averaged across conditions and subjects. Since this experiment is the first application of the virtual T-maze task while recording MEG, channels for statistical analysis were identified based on an exploratory approach. For each subject the channel carrying the peak RPT was extracted. Bar plots with black labels mark channels that were picked at least for one subject. The 2D topoplot on the right shows the locations of these channels identified as peak channels. **B**: Spectrograms with evoked power change relative to baseline (expressed as ratio) from -100 ms to 600 ms post-feedback separated by alleys (left alleys vs. right alleys) at EEG channel PO7. Raincloud plots show the distributions of theta power peaks and theta power peak latencies by alley (left in pink, right in blue) for PO7. Cyan dots mark the mean value of each distribution. Red and blue dots represent individual subject values. \**ns: not significant*

**Figure 5-figure supplement 1:**
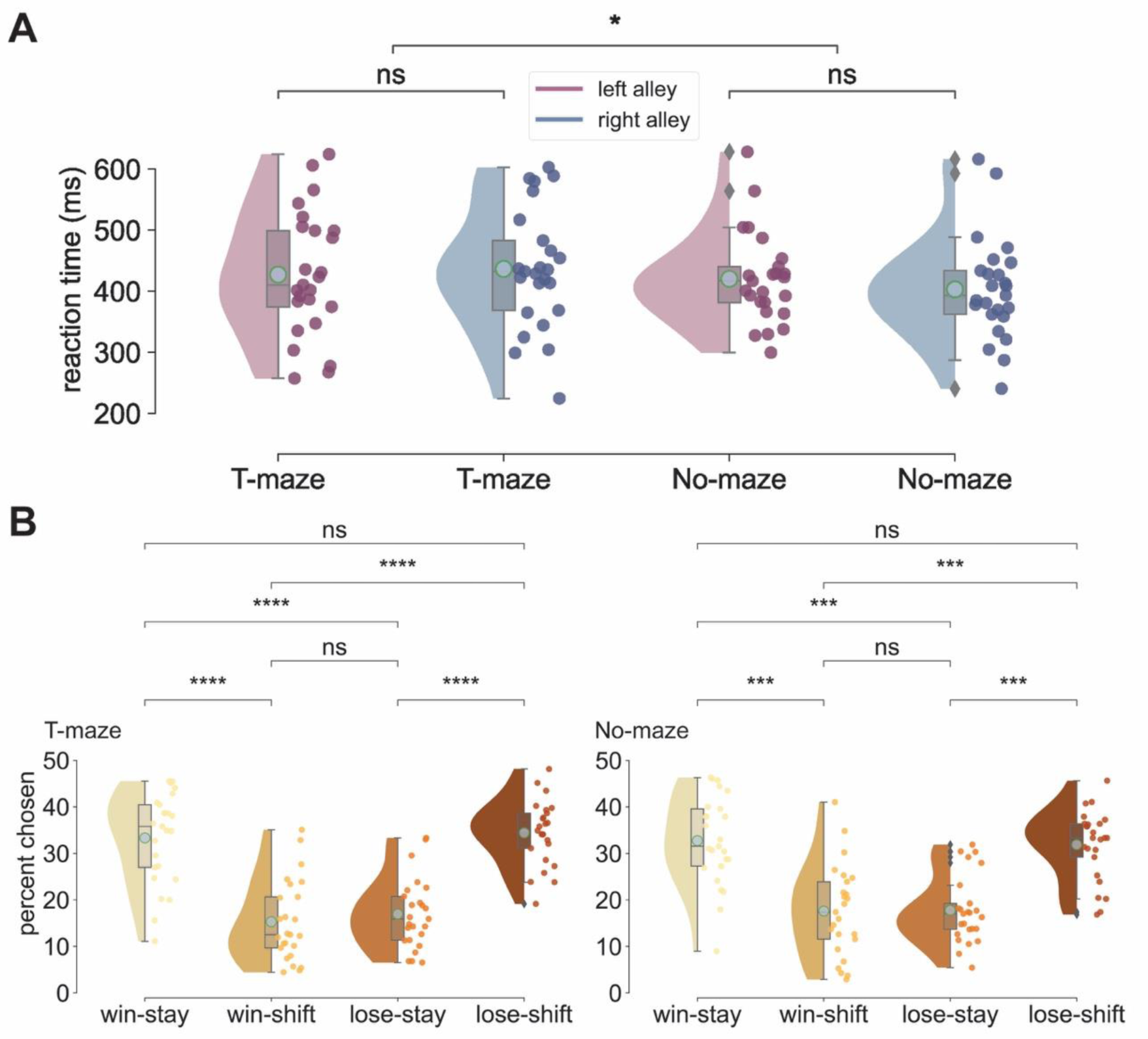
Experiment 2: Behavioral data analysis. **A**: Raincloud plots depicting the distribution of individual subject averages in milliseconds (ms) separated by alley (left in pink, right in blue) and condition (T-maze vs. No-maze) with cyan dots indicating the mean value of each distribution. A two-way repeated measures ANOVA run on the sum of left and right choices yielded neither a significant main effect of Feedback Alley Type (*F*(1, 24) = 0.478, *p* = 0.495, *η*^2^ = 0.019) nor an interaction between Feedback Alley Type and Task Type (*F*(1, 24) = 0.042, *p* = 0.837, *η*^2^ = 0.001), indicating that right and left alleys were chosen an equally in both the T-maze and No-maze condition. Similarly, there was no significant main effect of Feedback Alley Type when calculating the same ANOVA for the dependent variable reaction times, *F*(1, 24) = 0.453, *p* = 0.507, *η*^2^ = 0.018. Thus, left and right alleys were equivalent in reaction times. However, in the T-maze condition (M = 432 ms, SD = 95 ms) subjects responded significantly faster than in the No-maze condition (M = 412 ms, SD = 75 ms), as revealed by a significant main effect of the factor Task Type, *F*(1, 24) = 4.37, *p* = 0.047, *η*^2^ = 0.154. **B**: Percentage of response strategies relative to total choices. Same as in A, cyan dots are the mean of each distribution and dots colored like the density plots are each individual subject’s mean value. Consistent with previous studies, win-stay (M = 33%, SD = 10%) and lose-shift (M = 33%, SD = 7%) emerged as the most common response strategies. This was confirmed by a significant main effect of the factor response strategy (*F*(1, 24) = 27.24, *p* = 1.04 x 10^-7^, *η*^2^ = 0.531) in a two-way repeated measures ANOVA, testing the effects of the factors response strategy (win-stay, win-shift, lose-stay, lose-shift) and Task Type. Post-hoc tests (Bonferroni-Holm corrected) revealed that specifically win-stay was more frequent than win-shift (*t*(24) = 4.86, *p* = 8.78 x 10^-5^) and lose-stay (*t*(24) = 6.28, *p* = 5.18 x 10^-6^), and that lose-shift was more frequent than win-shift (*t*(24) = 6.37, *p* = 5.18 x 10^-6^) and lose-stay (*t*(24) = 5.85, *p* = 9.87 x 10^-6^). *ns: not significant, ***p < 0.0001, ****p < 0.00001*

**Figure 6-figure supplement 1:**
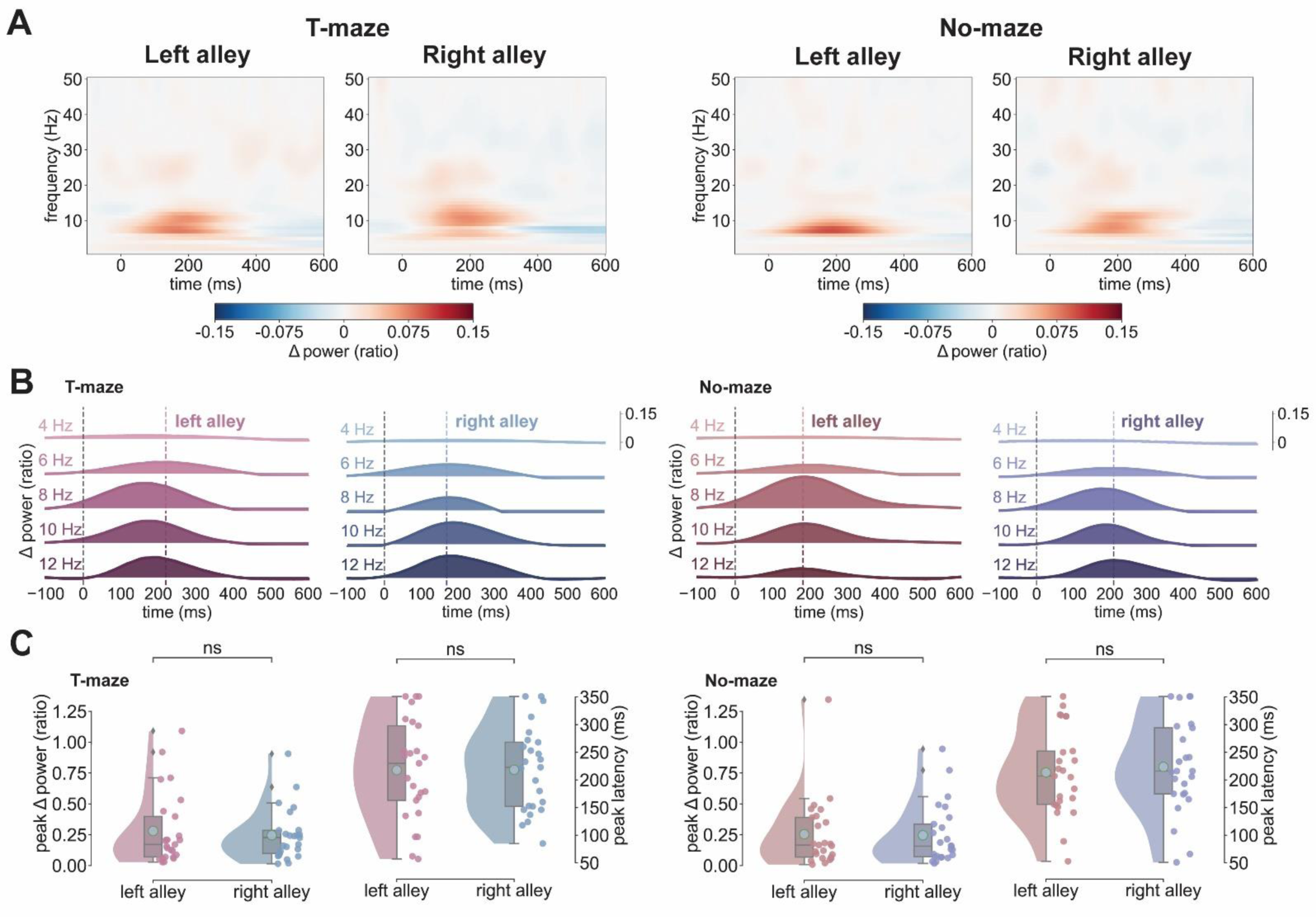
Left hemisphere RPT results. **A**: Spectrograms depicting the evoked power from -100 ms to 600 ms post-feedback at E107 (left hemisphere) separately for Task Type (T-maze: left panels, No- maze: right panels) and alley (left alleys vs. right alleys). Unlike on the right hemisphere, the two-way repeated measures ANOVA run for RPT peak latency with the factors Task (T-maze vs. No- maze) and Feedback Alley Type (left vs. right alley) neither yielded a significant interaction effect (*F*(1, 24) = 0.113, *p* = 0.739, *η*^2^ = 0.005) nor any significant main effects. Left alleys were followed by RPT peak latencies (M = 218 ms, SD = 89 ms) as right alleys (M = 218 ms, SD = 75 ms) in the T-maze and in the No-maze (left: M = 214 ms, SD = 77 ms; right: M = 224 ms, SD = 82 ms). The same was observed for the ANOVA run on RPT peak amplitudes. **B**: Ridge plots show the same results as in A divided into the delta band (1-4 Hz) and theta bands (5-6 Hz, 7-8 Hz, 9-10 Hz, and 11-12 Hz), with the average RPT peak timing for each condition marked by a dashed colored line. Results for the T-maze condition are on the left side and for the No-maze condition on the right side. **C**: Raincloud plots depict the RPT power peaks and peak latencies for the T-maze condition on the left and the No-maze condition on the right, with cyan dots showing the mean value of each distribution. Alley is shown on the x-axis and colored dots represent individual subject values. ns: not significant

**Figure 9-figure supplement 1:**
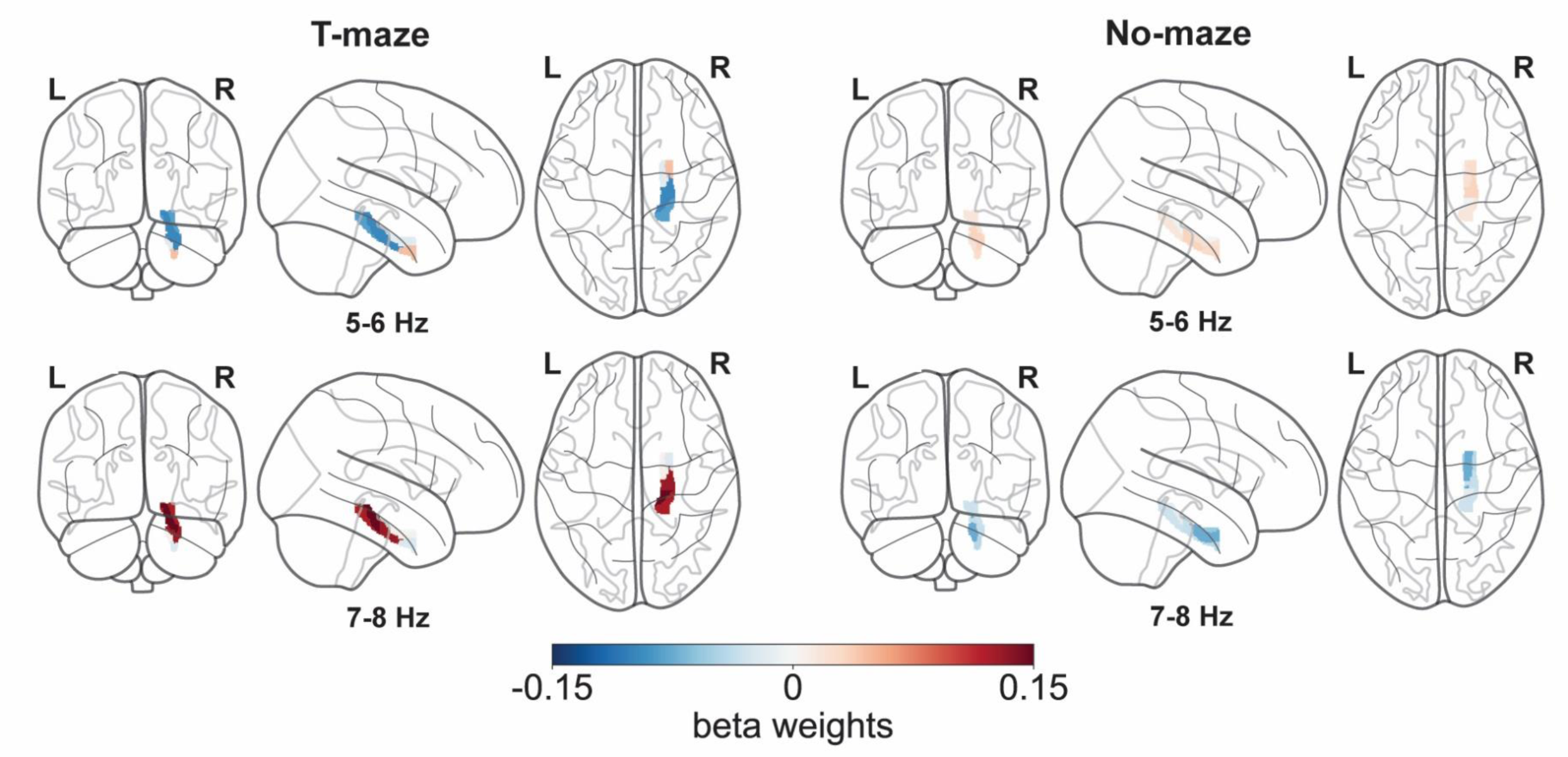
Right PHG beta weights. Glass brain plots show the 5-6 Hz and 7-8 Hz beta weights color-coded on their respective PHG ROIs (L: left hemisphere, R: right hemisphere) separated by T-maze (left) and No-maze condition (right). Color denotes the magnitude of beta weights from the respective LME model.

**Table 2-table supplement 1:**
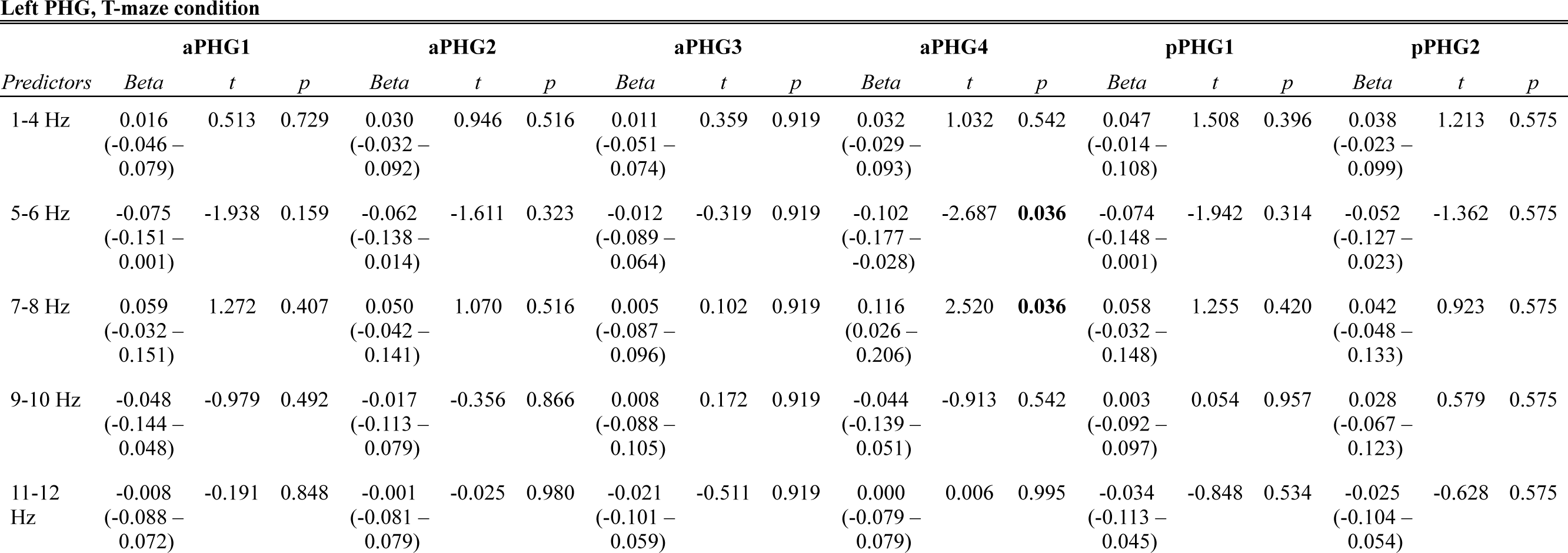
Fixed Effects table of the LME model run on single-trial left PHG ROI regression weights (columns) recorded during the T-maze condition. aPHG4 BOLD was significantly related to trial-by-trial variation in RPT. Increases in 5-6 Hz and 11-12 Hz power were associated with decreases in aPHG4 activation, while 7-8 Hz power was coupled to increases in aPHG4 activation.

**Table 2-table supplement 2:** Fixed Effects table of the LME models run on single-trial BOLD from the right PHG ROIs (columns) recorded during the No-maze condition.

**Right PHG, No-maze condition**
| *Predictors* | **aPHG1** |  |  | **aPHG2** |  |  | **aPHG3** |  |  | **aPHG4** |  |  | **pPHG1** |  |  | **pPHG2** |  |  |
| --- | --- | --- | --- | --- | --- | --- | --- | --- | --- | --- | --- | --- | --- | --- | --- | --- | --- | --- |
|  | *Beta* | *t* | *p* | *Beta* | *t* | *p* | *Beta* | *t* | *p* | *Beta* | *t* | *p* | *Beta* | *t* | *p* | *Beta* | *t* | *p* |
| 1-4 Hz | -0.035<br>(-0.093 –<br>0.024) | -1.168 | 0.365 | -0.066<br>(-0.125 –<br>-0.008) | -2.215 | 0.162 | -0.053<br>(-0.113 –<br>0.006) | -1.751 | 0.481 | -0.071<br>(-0.131 –<br>-0.011) | -2.330 | 0.120 | -0.042<br>(-0.102 –<br>0.017) | -1.392 | 0.487 | -0.032<br>(-0.092 –<br>0.028) | -1.052 | 0.904 |
| 5-6 Hz | -0.002<br>(-0.066 –<br>0.062) | -0.050 | 0.960 | 0.032<br>(-0.033 –<br>0.096) | 0.966 | 0.517 | 0.026<br>(-0.039 –<br>0.091) | 0.779 | 0.873 | 0.021<br>(-0.044 –<br>0.087) | 0.640 | 0.778 | 0.016<br>(-0.050 –<br>0.081) | 0.467 | 0.769 | -0.004<br>(-0.070 –<br>0.062) | -0.120 | 0.904 |
| 7-8 Hz | -0.056<br>(-0.128 –<br>0.015) | -1.540 | 0.365 | -0.035<br>(-0.107 –<br>0.037) | -0.945 | 0.517 | -0.042<br>(-0.115 –<br>0.031) | -1.120 | 0.789 | -0.075<br>(-0.148 –<br>-0.002) | -2.003 | 0.136 | -0.031<br>(-0.105 –<br>0.042) | -0.835 | 0.606 | -0.014<br>(-0.087 –<br>0.060) | -0.372 | 0.904 |
| 9-10 Hz | -0.059<br>(-0.140 –<br>0.022) | -1.419 | 0.365 | -0.042<br>(-0.124 –<br>0.039) | -1.021 | 0.517 | 0.004<br>(-0.079 –<br>0.087) | 0.103 | 0.932 | -0.012<br>(-0.095 –<br>0.071) | -0.282 | 0.778 | -0.049<br>(-0.132 –<br>0.034) | -1.167 | 0.487 | -0.039<br>(-0.123 –<br>0.044) | -0.930 | 0.904 |
| 11-12<br>Hz | 0.038<br>(-0.035 –<br>0.112) | 1.023 | 0.368 | 0.011<br>(-0.063 –<br>0.085) | 0.294 | 0.829 | -0.014<br>(-0.089 –<br>0.061) | -0.377 | 0.932 | -0.015<br>(-0.090 –<br>0.060) | -0.392 | 0.778 | 0.001<br>(-0.074 –<br>0.076) | 0.035 | 0.972 | 0.022<br>(-0.053 –<br>0.097) | 0.569 | 0.904 |

**Table 2-table supplement 3:** Fixed Effects table of the LME models run on single-trial BOLD from the left PHG ROIs (columns) recorded during the No-maze condition.

**Left PHG, No-maze condition**
| *Predictors* | **aPHG1** |  |  | **aPHG2** |  |  | **aPHG3** |  |  | **aPHG4** |  |  | **pPHG1** |  |  | **pPHG2** |  |  |
| --- | --- | --- | --- | --- | --- | --- | --- | --- | --- | --- | --- | --- | --- | --- | --- | --- | --- | --- |
|  | *Beta* | *t* | *p* | *Beta* | *t* | *p* | *Beta* | *t* | *p* | *Beta* | *t* | *p* | *Beta* | *t* | *p* | *Beta* | *t* | *p* |
| 1-4 Hz | -0.059<br>(-0.118 –<br>0.001) | -1.935 | 0.160 | -0.059<br>(-0.118 –<br>0.001) | -1.940 | 0.158 | 0.009<br>(-0.052 –<br>0.071) | 0.298 | 0.789 | -0.042<br>(-0.102 –<br>0.017) | -1.404 | 0.558 | -0.036<br>(-0.095 –<br>0.024) | -1.184 | 0.422 | -0.054<br>(-0.113 –<br>0.006) | -1.771 | 0.461 |
| 5-6 Hz | -0.017<br>(-0.083 –<br>0.048) | -0.519 | 0.850 | 0.003<br>(-0.062 –<br>0.068) | 0.084 | 0.933 | 0.025<br>(-0.043 –<br>0.092) | 0.715 | 0.789 | 0.013<br>(-0.051 –<br>0.078) | 0.403 | 0.687 | 0.034<br>(-0.031 –<br>0.100) | 1.039 | 0.422 | 0.036<br>(-0.029 –<br>0.102) | 1.094 | 0.548 |
| 7-8 Hz | -0.065<br>(-0.138 –<br>0.008) | -1.741 | 0.164 | -0.076<br>(-0.149 –<br>-0.004) | -2.060 | 0.158 | -0.056<br>(-0.131 –<br>0.019) | -1.471 | 0.789 | -0.046<br>(-0.118 –<br>0.027) | -1.232 | 0.558 | -0.040<br>(-0.113 –<br>0.033) | -1.077 | 0.422 | -0.048<br>(-0.121 –<br>0.025) | -1.286 | 0.548 |
| 9-10 Hz | -0.016<br>(-0.098 –<br>0.067) | -0.374 | 0.850 | -0.016<br>(-0.098 –<br>0.067) | -0.374 | 0.933 | 0.017<br>(-0.068 –<br>0.101) | 0.383 | 0.789 | -0.042<br>(-0.124 –<br>0.041) | -0.993 | 0.558 | -0.039<br>(-0.122 –<br>0.043) | -0.931 | 0.422 | -0.013<br>(-0.095 –<br>0.070) | -0.304 | 0.913 |
| 11-12<br>Hz | 0.007<br>(-0.068 –<br>0.081) | 0.177 | 0.859 | 0.008<br>(-0.067 –<br>0.082) | 0.200 | 0.933 | -0.010<br>(-0.087 –<br>0.066) | -0.268 | 0.789 | 0.034<br>(-0.040 –<br>0.108) | 0.893 | 0.558 | -0.005<br>(-0.079 –<br>0.070) | -0.129 | 0.898 | 0.002<br>(-0.072 –<br>0.077) | 0.064 | 0.949 |

**Table 2-table supplement 4:** Fixed Effects table of the LME models run on single-trial BOLD from the control ROIs (left and right Heschl’s Gyrus, Precuneus; columns) recorded during the T-maze condition.

**Control ROIs, T-maze condition**
| *Predictors* | **Left Heschl's Gyrus** |  |  | **Right Heschl's Gyrus** |  |  | **Precuneus** |  |  |
| --- | --- | --- | --- | --- | --- | --- | --- | --- | --- |
|  | *Estimates* | *CI* | *p* | *Estimates* | *CI* | *p* | *Estimates* | *CI* | *p* |
| 1-4 Hz | -0.015 | -0.069 – 0.038 | 0.924 | -0.019 | -0.071 – 0.033 | 0.716 | -0.004 | -0.084 – 0.075 | 0.916 |
| 5-6 Hz | -0.001 | -0.046 – 0.044 | 0.950 | -0.005 | -0.048 – 0.039 | 0.833 | -0.041 | -0.107 – 0.025 | 0.446 |
| 7-8 Hz | 0.010 | -0.030 – 0.050 | 0.924 | 0.028 | -0.011 – 0.067 | 0.458 | 0.027 | -0.032 – 0.086 | 0.560 |
| 9-10 Hz | -0.003 | -0.052 – 0.047 | 0.950 | -0.008 | -0.056 – 0.040 | 0.833 | 0.055 | -0.017 – 0.128 | 0.400 |
| 11-12 Hz | -0.013 | -0.059 – 0.033 | 0.924 | -0.018 | -0.062 – 0.027 | 0.716 | -0.085 | -0.152 – -0.018 | 0.081 |

**Table 2-table supplement 5.** : Fixed Effects table of the LME models run on single-trial BOLD from the Glasser atlas ROIs (columns) recorded during the T-maze condition. Only the parietal-temporal ROIs with at least one significant beta weight are shown.

**Parietal-temporal Glassor ROIs, T-maze condition**
| *Predictors* | Parahippocampal Area 3 (right) |  |  | Perirhinal Ectorhinal Cortex (right) |  |  | Ventral Intraparietal Complex (left) |  |  |
| --- | --- | --- | --- | --- | --- | --- | --- | --- | --- |
|  | *Beta* | *t* | *p* | *Beta* | *t* | *p* | *Beta* | *t* | *p* |
| 1-4 Hz | -0.006<br>(-0.068 – 0.055) | -0.205 | 0.837 | -0.057<br>(-0.118 – 0.004) | -1.848 | 0.130 | 0.018<br>(-0.043 – 0.078) | 0.567 | 0.685 |
| 5-6 Hz | -0.097<br>(-0.172 – -0.022) | -2.543 | **0.045** | 0.109<br>(0.035 – 0.183) | 2.898 | **0.023** | -0.101<br>(-0.176 – -0.027) | -2.671 | **0.046** |
| 7-8 Hz | 0.112<br>(0.022 – 0.203) | 2.435 | **0.045** | 0.098<br>(0.008 – 0.187) | 2.149 | 0.096 | 0.076<br>(-0.014 – 0.166) | 1.656 | 0.196 |
| 9-10 Hz | -0.051<br>(-0.146 – 0.044) | -1.052 | 0.439 | 0.008<br>(-0.086 – 0.102) | 0.171 | 0.865 | -0.032<br>(-0.126 – 0.062) | -0.665 | 0.685 |
| 11-12 Hz | -0.013<br>(-0.092 – 0.066) | -0.315 | 0.837 | -0.027<br>(-0.105 – 0.052) | -0.667 | 0.606 | 0.000<br>(-0.078 – 0.079) | 0.006 | 0.995 |

**Table 2-table supplement 6:** Fixed Effects table of the LME models run on single-trial BOLD from the Glasser atlas ROIs (columns) recorded during the T-maze condition. Only the frontal ROIs with at least one significant beta weight are shown.

Frontal Glassor ROIs, T-maze condition
| *Predictors* | Anterior Area 24' (left) |  |  | Area 33' (left) |  |  | Dorsal Area 8A (left) |  |  | Ventral Area 8A (left) |  |  | Anterior Area IFJ (right) |  |  | Posterior Area IFJ (right) |  |  | Dorsal Area 32 (left) |  |  | Posterior Area 32' (left) |  |  |
| --- | --- | --- | --- | --- | --- | --- | --- | --- | --- | --- | --- | --- | --- | --- | --- | --- | --- | --- | --- | --- | --- | --- | --- | --- |
|  | *Beta* | *t* | *p* | *Beta* | *t* | *p* | *Beta* | *t* | *p* | *Beta* | *t* | *p* | *Beta* | *t* | *p* | *Beta* | *t* | *p* | *Beta* | *t* | *p* | *Beta* | *t* | *p* |
| 1-4 Hz | 0.029<br>(-0.032 –<br>0.090) | 0.935 | 0.420 | 0.024<br>(-0.034 –<br>0.082) | 0.818 | 0.620 | 0.011<br>(-0.050 –<br>0.072) | 0.359 | 0.841 | 0.016<br>(-0.045 –<br>0.077) | 0.519 | 0.604 | 0.027<br>(-0.031 –<br>0.085) | 0.916 | 0.432 | 0.007<br>(-0.051 –<br>0.065) | 0.225 | 0.840 | 0.010<br>(-0.050 –<br>0.069) | 0.323 | 0.747 | 0.012<br>(-0.049 –<br>0.072) | 0.375 | 0.849 |
| 5-6 Hz | -0.131<br>(-0.206 –<br>-0.057) | -3.471 | **0.003** | -0.089<br>(-0.160 –<br>-0.018) | -2.449 | **0.044** | -0.095<br>(-0.170 –<br>-0.020) | -2.500 | **0.025** | -0.131<br>(-0.205 –<br>-0.057) | -3.457 | **0.002** | -0.114<br>(-0.185 –<br>-0.043) | -3.139 | **0.010** | -0.069<br>(-0.140 –<br>0.002) | -1.913 | 0.112 | -0.090<br>(-0.163 –<br>-0.018) | -2.436 | **0.045** | -0.127<br>(-0.201 –<br>-0.052) | -3.350 | **0.005** |
| 7-8 Hz | 0.151<br>(0.061 –<br>0.241) | 3.295 | **0.003** | 0.112<br>(0.026 –<br>0.198) | 2.556 | **0.044** | 0.120<br>(0.030 –<br>0.210) | 2.609 | **0.025** | 0.179<br>(0.090 –<br>0.269) | 3.920 | **0.001** | 0.129<br>(0.043 –<br>0.215) | 2.937 | **0.010** | 0.136<br>(0.050 –<br>0.222) | 3.108 | **0.012** | 0.119<br>(0.031 –<br>0.207) | 2.651 | **0.045** | 0.131<br>(0.041 –<br>0.221) | 2.871 | **0.013** |
| 9-10 Hz | -0.065<br>(-0.159 –<br>0.029) | -1.358 | 0.262 | -0.040<br>(-0.130 –<br>0.050) | -0.871 | 0.620 | -0.059<br>(-0.154 –<br>0.035) | -1.228 | 0.330 | -0.081<br>(-0.175 –<br>0.013) | -1.683 | 0.139 | -0.067<br>(-0.158 –<br>0.023) | -1.470 | 0.213 | -0.064<br>(-0.154 –<br>0.026) | -1.402 | 0.242 | -0.074<br>(-0.166 –<br>0.018) | -1.577 | 0.230 | -0.004<br>(-0.097 –<br>0.090) | -0.074 | 0.941 |
| 11-12 Hz | -0.004<br>(-0.082 –<br>0.075) | -0.094 | 0.925 | 0.017<br>(-0.058 –<br>0.092) | 0.453 | 0.706 | 0.008<br>(-0.071 –<br>0.087) | 0.200 | 0.841 | 0.023<br>(-0.056 –<br>0.101) | 0.570 | 0.604 | -0.020<br>(-0.095 –<br>0.055) | -0.533 | 0.594 | 0.008<br>(-0.067 –<br>0.083) | 0.201 | 0.840 | 0.013<br>(-0.063 –<br>0.090) | 0.345 | 0.747 | -0.029<br>(-0.108 –<br>0.049) | -0.737 | 0.849 |

